# Antigen perception in T cells by long-term Erk and NFAT signaling dynamics

**DOI:** 10.1101/2023.06.01.543260

**Authors:** Matthew J Wither, William L White, Sriram Pendyala, Paul J Leanza, Doug Fowler, Hao Yuan Kueh

## Abstract

Immune system threat detection hinges on T cells’ ability to perceive varying peptide major-histocompatibility complex (pMHC) antigens. As the Erk and NFAT pathways link T cell receptor engagement to gene regulation, their signaling dynamics may convey information about pMHC inputs. To test this idea, we developed a dual reporter mouse strain and a quantitative imaging assay that, together, enable simultaneous monitoring of Erk and NFAT dynamics in live T cells over day-long timescales as they respond to varying pMHC inputs. Both pathways initially activate uniformly across various pMHC inputs, but diverge only over longer (9+ hrs) timescales, enabling independent encoding of pMHC affinity and dose. These late signaling dynamics are decoded via multiple temporal and combinatorial mechanisms to generate pMHC-specific transcriptional responses. Our findings underscore the importance of long timescale signaling dynamics in antigen perception, and establish a framework for understanding T cell responses under diverse contexts.

**SIGNIFICANCE STATEMENT:** To counter diverse pathogens, T cells mount distinct responses to varying peptide-major histocompatibility complex ligands (pMHCs). They perceive the affinity of pMHCs for the T cell receptor (TCR), which reflects its foreignness, as well as pMHC abundance. By tracking signaling responses in single living cells to different pMHCs, we find that T cells can independently perceive pMHC affinity vs dose, and encode this information through the dynamics of Erk and NFAT signaling pathways downstream of the TCR. These dynamics are jointly decoded by gene regulatory mechanisms to produce pMHC-specific activation responses. Our work reveals how T cells can elicit tailored functional responses to diverse threats and how dysregulation of these responses may lead to immune pathologies.

## INTRODUCTION

The immune system can mount tailored responses to different threats while sparing healthy tissues. This remarkable selectivity stems from T cells’ ability to obtain information about the threat from peptide antigens presented on major histocompatibility complex ligands (pMHCs)(1, 2). pMHCs that bind the T cell receptor (TCR) with high affinity derive exclusively from foreign sources, due to sharp affinity threshold for negative selection (3, 4). On the other hand, lower-affinity pMHCs can derive either from foreign sources or self. The dose of pMHCs presented to the T cell yields complementary information: high doses of pMHCs can convey pathogen virulence or infection severity, or simply reflect high abundance of self-antigen.

Despite the importance of antigen sensing for immunity, how T cells perceive pMHC affinity and dose to generate tailored functional responses remains incompletely understood. Early studies suggested that T cells respond to high affinity pMHCs while ignoring high doses of low affinity pMHCs (5–10). This selectivity was proposed to result from kinetic proofreading in the TCR signaling pathway (6, 11–19), which enables selective responses to pMHCs with long binding lifetimes. However, while there is compelling evidence for kinetic proofreading (20, 21), T cells do not ignore low-affinity pMHC ligands (22–24). More recent studies indicate that T cells enact a common gene activation program amid different pMHC affinities, albeit with varying kinetics (25, 26). pMHC differences can tune these gene programs (25, 26), thereby modulating downstream outcomes (27–30), with their dose and affinity exerting distinct regulatory effects (31–34). These findings suggest that the TCR signaling network can sense pMHC affinity and dose to translate this information into distinct functional outcomes.

As the Erk and NFAT signaling pathways are primary links between TCR engagement and the regulation of gene transcription (35), they are well-positioned to convey pMHC information in the cell. Upon pMHC engagement, the TCR activates a series of intracellular signaling molecules, leading to activation of both Erk and NFAT pathways. Both pathways activate within minutes upon antigen encounter, and over these short timescales, appear to do so in an all-or-one, “digital” manner, with activated cells showing similar signaling levels regardless of pMHC affinity or dose (6, 36–38). However, T cells can be exposed to pMHCs for up to 24 hrs after initial encounter *in vivo* (39, 40) through stable contacts with dendritic cells in lymph nodes. Over these longer timescales, it is unclear whether Erk and NFAT signaling remains uniform to varying pMHC inputs, or whether they may show variable responses to different pMHC inputs only upon prolonged exposure. Indeed, longer timescale signaling dynamics (∼hrs) underlies input encoding in other systems (41–43). However, its role in T cell antigen perception remain unclear, as most TCR signaling measurements, to our knowledge, have been performed over short timescales (<3 hrs).

NFAT and Erk signaling leads to nuclear accumulation and activation of the NFAT transcription factor (TF) and AP-1 family transcription factors (TFs) respectively (44). As these TFs can show different modes of cooperation by which they regulate their target genes, they could enable cells to translate pMHC-specific Erk and NFAT signaling activities to distinct transcriptional responses. NFAT and AP-1 can bind to composite DNA binding sites for both TFs cooperatively (45, 46), a mode of action that underlies regulation of many T cell activation and effector genes (*Irf4, Il2ra*). Disruption of NFAT:AP-1 impairs T cell effector responses and results in an inability to clear chronic infections and tumors (47). At the same time, NFAT can work independently from AP-1, a mode of regulation for exhaustion- or anergy-associated genes (e.g. *Pdcd1*) (45, 48–50). These different *cis*-regulatory mechanisms may allow T cells to generate distinct genomic programs to different pMHC inputs.

In this study, we investigated the role for Erk and NFAT signaling dynamics in antigen perception, specifically seeking to measure signaling pathway dynamics in living T cells over daylong timescales over which T cells contact antigen-presenting cells, and over which the gene regulatory responses unfold (39). To do so, we developed a dual-pathway fluorescent reporter mouse strain that concurrently reveals Erk and NFAT signaling activity in the same cell. We then developed an *in vitro* imaging assay to quantify the first 30 hrs of Erk and NFAT signaling dynamics in CD8^+^ T cells responding to alterations in pMHC affinity and dose. We then use RNA-sequencing to investigate how these dynamics generate specific affinity- and dose-dependent gene programs and propose a model to explain how diverse input-dependent gene expression arises from these signaling activities.

## RESULTS

### Dual-pathway reporter mouse enables continuous measurements of Erk and NFAT activity in living T cells

To quantify Erk and NFAT signaling dynamics in living T cells, we developed a reporter system to concurrently measure the activities of both pathways in live CD8^+^ T cells (Fig. 1A). This system consists of (1) The N-terminal regulatory domain of mouse NFATc2 (residues 1-399) fused to mRuby3 (38, 51, 52); (2) An Erk kinase-translocation reporter (Erk-KTR) (53) fused to BFP; and (3) Histone 2B (H2B) fused to iRFP for nuclear segmentation. We generated a “dual-pathway reporter” (DPR) mouse strain with this reporter cassette inserted into the Rosa26 locus downstream of a loxP-flanked STOP codon (*Rosa26^LSL-DPR^*). *Rosa26^LSL-DPR/LSL-DPR^*mice were crossed to the P14 transgenic TCR strain to enable stimulation with altered peptide ligands (APLs) for the cognate gp_33-41_ peptide from LCMV complexed to H-2D^b^ MHC-1 (54). DPR^+^/P14^+^ mice were then crossed to a Cre-ERT2 strain and treated with tamoxifen for reporter induction (Fig. S1A).

**Figure 1.**
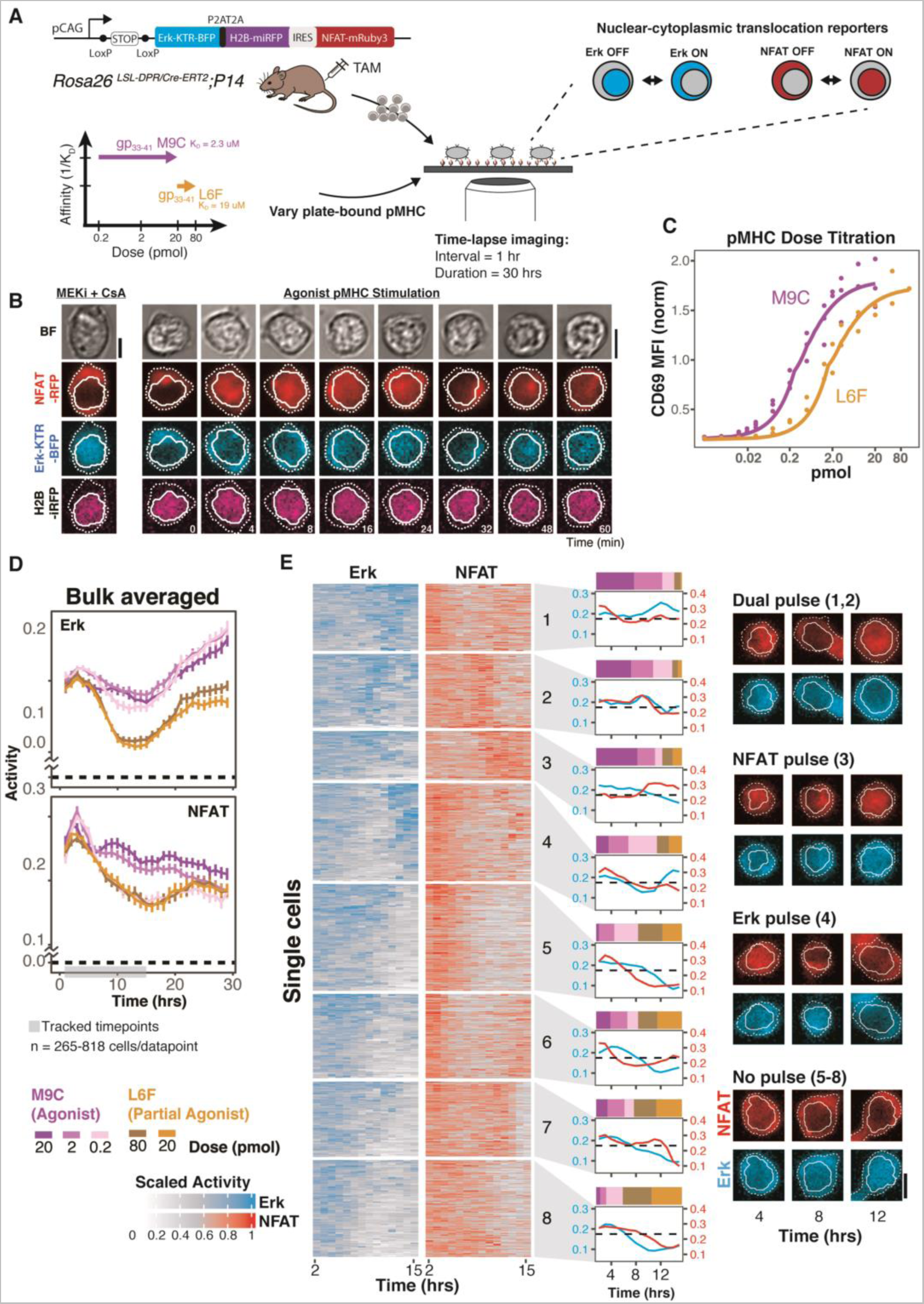
Erk and NFAT signaling varies with pMHC affinity and dose over long timescales. (A) Schematic of the DPR mouse strain used to quantify Erk and NFAT signaling dynamics in live T cells. The NFAT reporter translocates from the cytoplasm to the nucleus when active. Erk-KTR translocates from the nucleus to cytoplasm when active. To assess pMHC-dependent signaling, we used various doses of two APLs that bind the P14 TCR with ∼8 fold difference in binding affinity. (B) Representative cell images of reporter T cells responding to a saturating dose of agonist pMHC over the first hour of stimulation, compared to a negative control cell treated with NFAT (Cyclosporin A, CsA) and Erk (Trametinib, MEKi) inhibitors. Nuclear and cytoplasmic segmentation masks are shown in solid and dashed lines, respectively. Scale bars = 5 um. (C) P14 Jurkat T cell dose-response curve based on CD69 upregulation 4 hrs after stimulation with each of the two APLs. Y-axis is MFI normalized to the EC_50_ for each APL. (D) Mean +/- 99% confidence intervals for Erk and NFAT activity over the first 30 hrs of stimulation with each of the five pMHC conditions from ∼180k cell images from two independent replicates with 265-818 cells (∼600 average) per data point. (E) 220 single cell traces per condition from the data in (D) were clustered on their scaled dual-pathway signaling activities from 2-15 hrs, and partitioned into 8 clusters. The mean unscaled response, and the proportion of cells from each pMHC condition for each cluster is shown to the right of the heatmap, with dashed lines indicating the mean Erk and NFAT level across all cells in the analysis. On the far right are representative cell images for four broad types of combinatorial Erk and NFAT responses classified from the eight clusters indicated in parentheses. Nuclear and cytoplasmic segmentations are shown in solid and dashed lines, respectively. Scale bar = 5 um.

To measure Erk and NFAT dynamics in DPR^+^/P14 (reporter) CD8^+^ T cells responding to pMHCs of different doses and affinities, we developed a quantitative imaging assay where T cells are cultured on purified, surface-immobilized pMHCs (55, 56) to ensure they experience uniform, defined inputs and then imaged at regular intervals with time-lapse microscopy (Fig. 1A). We stimulated reporter T cells with a high affinity gp_33-41_ pMHC variant (M9C, K_D_ = 2.3 μM) (57), imaged them at four-minute intervals for one hour, and quantified reporter localization (Fig. 1B and S1B). Both Erk and NFAT activated within minutes (Fig. S1C-E), consistent with their rapid activation upon TCR engagement (36, 38), and remained active for the duration of the observation. Activation was abolished by MEK (Trametinib; MEKi) and Calcineurin (Cyclosporin A; CsA) inhibitor treatment to block activation of Erk and NFAT, respectively (Fig. 1B), providing a baseline for reporter inactivity.

### T cells perceive pMHC affinity and dose using long-term Erk and NFAT signaling dynamics

Both Erk and NFAT pathways activate in a digital, all-or-none manner within the first hour of exposure to varying pMHC inputs (36, 58). However, T cells can form stable contacts with dendritic cells for up to 24 hrs after initial contact (39, 40), and may respond divergently to differing pMHC affinity and dose only over these longer timescales. To test this, we measured Erk and NFAT signaling activity in T cells over 30 hrs of stimulation with five pMHC conditions that vary in affinity or dose. We selected pMHC conditions based on a titration of the high affinity agonist, gp33-M9C, and the lower affinity partial agonist, gp33-L6F (K_D_ = 19 uM) (57) (Fig. 1A), and the resulting expression of the T cell activation marker CD69 (3) (Fig. 1C and S2B-C). To assess T cells’ ability to sense pMHC affinity, we chose saturating doses of each pMHC (2 and 20 pmol M9C vs. 20 and 80 pmol L6F), which yield equivalent CD69 activation potencies (3). To assess T cells’ ability to sense pMHC dose, we chose a dose of M9C (0.2 pmol) just below that needed for half-maximal CD69 activation. We then stimulated reporter CD8^+^ T cells with each of these five pMHC conditions and acquired images at 1 hr intervals for 30 hours (Supplementary Videos 1-5). We quantified the average Erk and NFAT activities in single cells (Fig. 1D), and extracted Erk and NFAT activity traces for ∼1,500 cells over the first 15 hrs (Fig. 1E).

Upon stimulation by different pMHC inputs, both pathways activated, as expected, and reached similar maximal levels of activity within the first hour after pMHC encounter (Fig. 1D and S1D). Lowering pMHC dose by a hundred-fold did not reduce maximal Erk and NFAT activity levels but reduced the rate of NFAT activation (Fig. S1E), consistent with prior studies showing that signal strength modulates the speed of all-or-none T cell activation (25, 26). However, after several hours of continued stimulation, divergent signaling dynamics emerged in a pMHC-dependent manner (Fig. 1D-E). Mean Erk activity dropped after 8-20 hrs of stimulation with L6F, but remained high for M9C at both high and low doses. In contrast, mean NFAT activity dropped rapidly for all doses of L6F, as well as for low dose of M9C, but remained persistently higher with higher doses of M9C. These signaling differences were not due to differences in pMHC stability, as both L6F and M9C were found to show negligible reduction in activity over these timescales when plate-immobilized (Fig. S2A). Furthermore, similar pMHC-dependent differences were also observed in non-reporter T cells immunostained for endogenous ppErk or nuclear-localized NFAT1 (Fig. S2D), indicating that our reporter system faithfully captures endogenous Erk/NFAT signaling states in T cells.

We next determined the dynamic features of Erk and NFAT signaling in single cells that encode information about pMHC affinity and dose. To do so, we performed hierarchical clustering on 1,100 single cell signaling responses from the 5 different pMHC conditions and identified eight clusters that showed distinct long-term signaling dynamics (Fig. 1E). While some cell clusters showed a monotonic decay in the activity of both pathways (clusters 5-8), others showed a second pulse of activity from either NFAT (cluster 3), Erk (cluster 4) or from both pathways (clusters 1-2). Different clusters were enriched for cells stimulated with different pMHC inputs, suggesting a role for these pulsed dynamics in encoding pMHC affinity and dose information. Thus, we quantified the incidence and dynamic features of Erk and NFAT pulsing in single cells and analyzed how the dynamics vary across different pMHC conditions (Fig. 2A-E).

**Figure 2.**
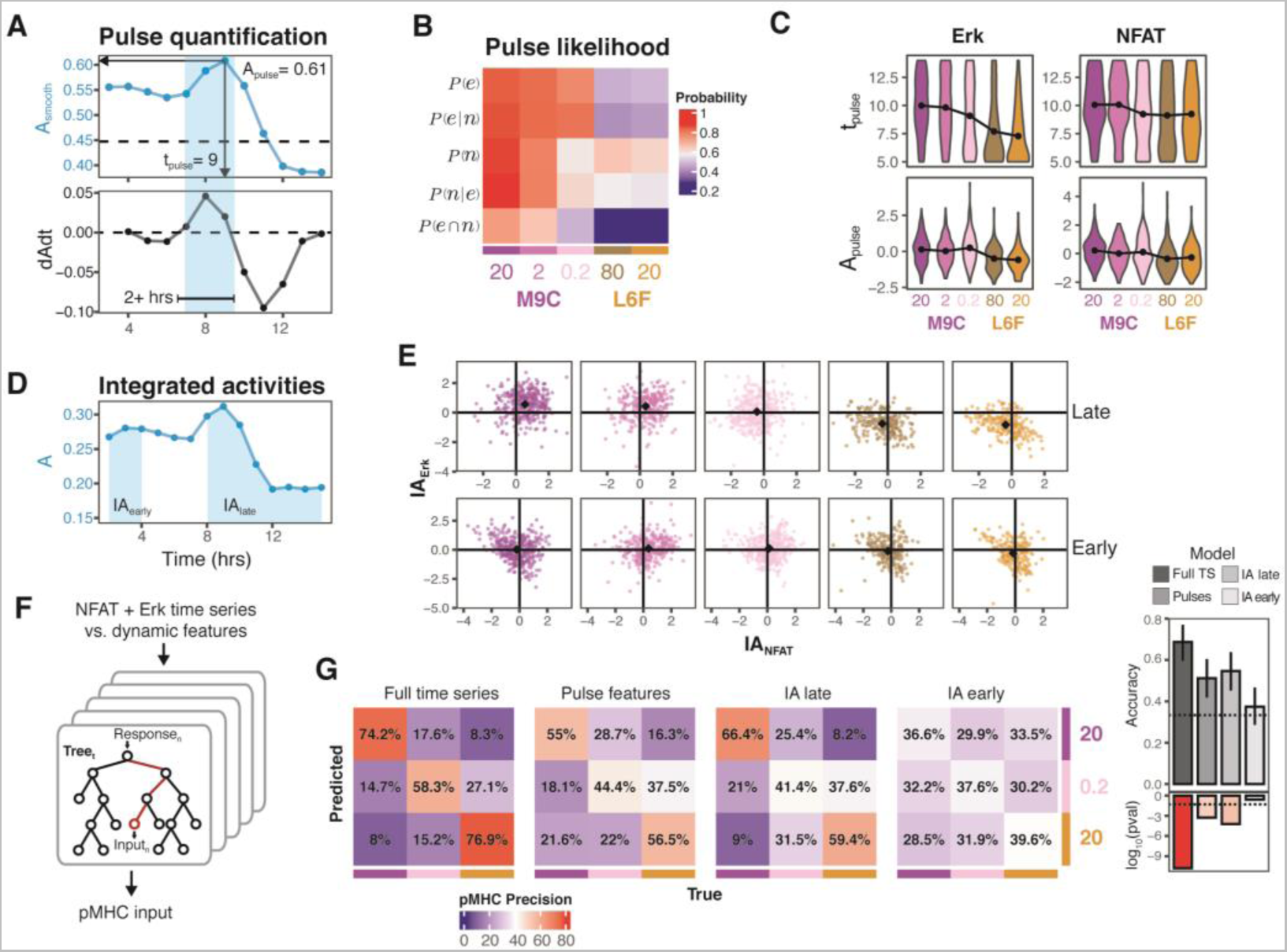
T cells independently encode pMHC affinity and dose information through late Erk and NFAT pulsatile dynamics. (A) Quantification of pulsatile dynamics from single cell traces. (B) Probabilities of NFAT and/or Erk pulses occurring for each pMHC condition. (C) Distributions of pulse time and scaled amplitude by pMHC condition, gated on cells with a pulse for the respective pathway. Black connected dots represent distribution means. (D) Quantification of early and late integrated activities from single cells. (E) 2D scatter plots of scaled early and late integrated activities for Erk and NFAT in each cell, separated by pMHC condition with colors matching those in (C). Black diamond is the mean for each condition. n = 1,510 total cells. (F) A random forest machine learning classification model was used to make predictions on the input pMHC condition given a dual-pathway signaling response in single cells. Four different random forest models were trained with different signaling predictors including: Full time series, pulse features (t_pulse_ + A_pulse_), IA_late_, and IA_early_ for both pathways. (G) Random forest model precision in predicting the input pMHC affinity and dose regime (high dose, high affinity; low dose, high affinity; high dose, low affinity) for each of the four sets of signaling predictors. Accuracy of each model is quantified on the right with the log_10_(p-values) beneath (dashed line is p-value = 0.05), testing if the model accuracy is greater than that by random chance (Accuracy = 0.33). Error bars represent 95% confidence intervals obtained from an exact binomial test.

We found that high doses of M9C elicited a second Erk and NFAT pulse with high probability, whereas equivalently high doses of L6F elicited substantially lower probabilities of generating second Erk and NFAT pulses (Fig. 2B). Strikingly, lowering the dose of M9C had a distinct effect from using the lower-affinity L6F at an equivalent dose, selectively reducing the second NFAT pulse probability while maintaining high Erk pulse probabilities. We also observed pMHC-dependent effects on pulse dynamics, such that even when L6F stimulation elicited second pulses, they had reduced amplitude and earlier onset compared to those elicited with M9C (Fig. 2B and C). We observed the same trends when using early and late integrated activities as an alternate measure of the pulsatile activity (Fig 2E). Taken together, these results indicate that Erk pulsing encodes information about pMHC affinity and NFAT pulsing encodes information about pMHC dose.

To quantify the extent to which late NFAT and Erk pulsing dynamics enables discrimination between different pMHC inputs, we trained a random forest classification model to distinguish between three pMHC dose and affinity regimes using quantified pulse dynamics (“pulse features”) (Fig. 2F-G and Fig. S2F). As a comparison, we also trained this model to use either the full time series for both pathways, or their early or late integrated activities. We found that the model trained on the full time series responses can discriminate between the three pMHC regimes with reasonable precision (∼75%). Additionally, while early integrated activity provided little discriminatory power (∼33% precision), as expected (Fig. 2G), long-term pulsing dynamics (∼55% precision) and integrated activity (∼60% precision) also enabled discrimination of the three input regimes, allowing for the independent encoding of pMHC dose and affinity though NFAT and Erk signaling, respectively (Fig. S2G).

### T cell activation genes exhibit distinct modes of decoding Erk and NFAT signaling

Genes downstream of TCR signaling integrate AP-1 and NFAT inputs in distinct ways. Therefore, tuning the activation of each TF through the differential long-term Erk and NFAT dynamics we discovered could plausibly lead to distinct gene expression programs. To explore this possibility, we used RNA-sequencing to identify gene programs that depend on Erk and NFAT signaling at long timescales. We subjected CD8^+^ T cells to strong stimulation (2 pmol M9C) to elicit maximal signaling, then treated cells with either MEKi and/or CsA after 9 hrs (Fig. 3A), when signaling activities diverge for different pMHC inputs (Fig. 1). Live imaging confirmed signaling inhibition as expected t(Fig. 3B). Cells were then subjected to bulk RNA sequencing after 30 hrs of stimulation and analyzed for differential gene expression to identify targets of late Erk and/or NFAT signaling. This analysis identified 1,089 differentially expressed genes (DEGs) regulated by Erk and/or NFAT signaling, with substantial overlap between their target genes (Fig. S3B), consistent with the cooperativity of NFAT and AP-1. We classified each gene as being strongly (s) or weakly (w) regulated based on their magnitude of expression fold change upon signaling inhibition (Fig. 3C).

**Figure 3.**
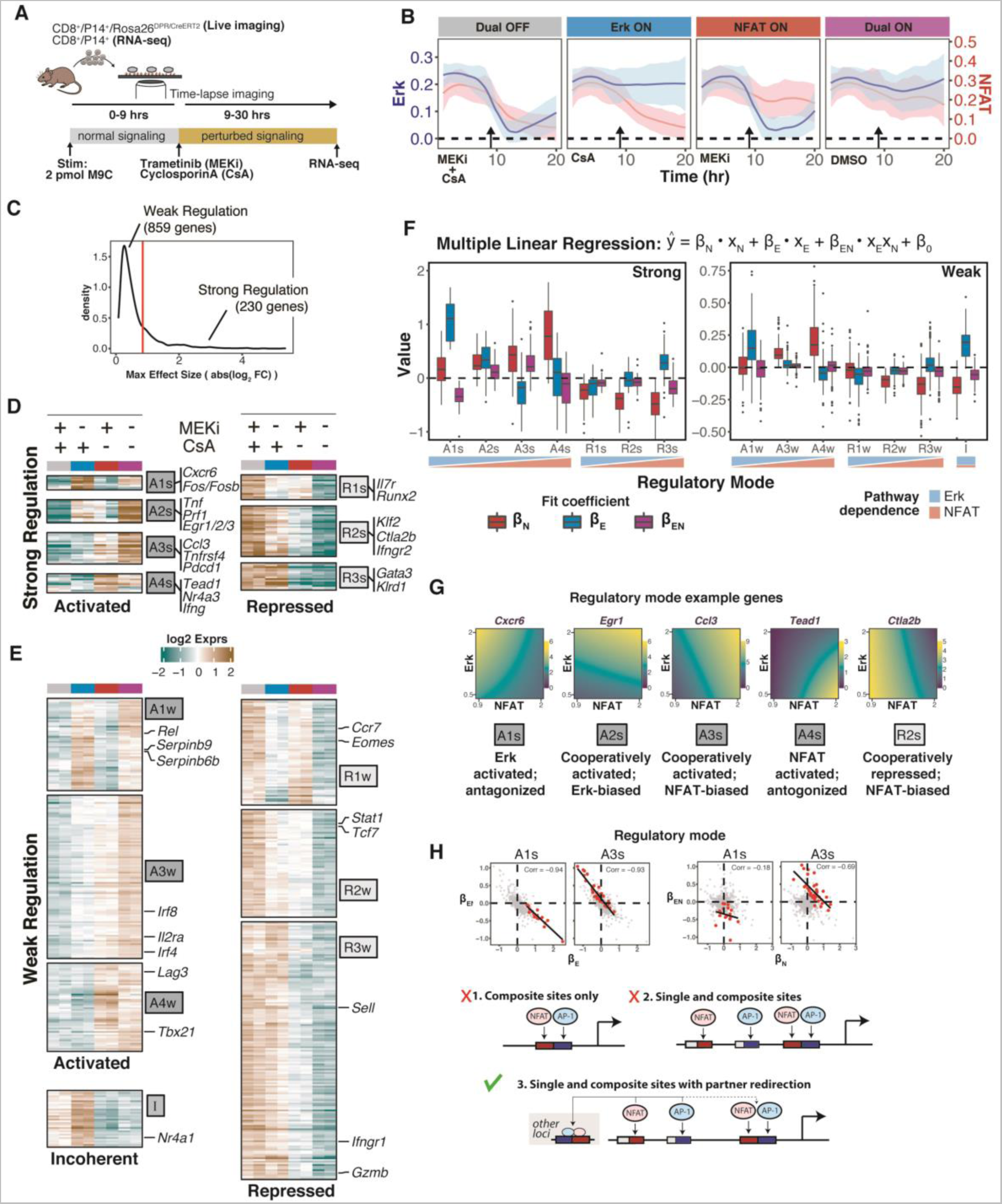
Target genes of Erk and NFAT show distinct modes for decoding signaling activity. (A) Live imaging and bulk RNA sequencing of CD8^+^ T cells treated MEKi and/or CsA. (B) Mean +/- SD from 96-110 single cell Erk and NFAT activity traces in response to each perturbation. (C) DEGs from bulk RNA-seq were identified based on a q-value threshold of 0.001 and classified as being strongly or weakly regulated by NFAT and/or Erk relative to a maximum fold change from any treatment of 1.8. (D-E) Hierarchical clustering of genes strongly (D) and weakly (E) regulated by Erk and NFAT signaling. Classification of regulatory modes was determined by distinct patterns of expression change due to inhibitor treatments. Genes of interest are identified from each regulatory mode. “A” = Activated, “R” = Repressed, “I” = Incoherent, “s” = strong, “w” = weak. (F) Boxplots of β coefficients from multiple linear regression models and grouped by regulatory mode. (G) Characterization of five regulatory modes. Example genes for each mode are shown with the model prediction of their expression with changing levels of Erk and NFAT signaling. (H) Representative pairwise correlations between β_EN_ vs β_E_ and β_EN_ vs β_N_ for the A1s and A3s genes. All DEGs are shown in gray dots with genes of each regulatory mode shown in red. Linear regressions are shown in black. Pearson correlation coefficients are indicated. *trans* TF partner redirection is required to reproduce the observed β coefficient correlations between single and combinatorial signaling sensitivities.

Analysis of strongly- and weakly-regulated genes using hierarchical clustering and linear regression (Fig. 3D-F and S3C) revealed a spectrum of regulatory modes by which distinct clusters of genes differentially integrate Erk and NFAT signaling inputs. Up-regulated genes could be either primarily activated by Erk (A1s, A1w, A2s, A2w) or by NFAT (A3s, A3w, A4s, A4w) (Fig. 3D-E), with these two subsets having highest sensitivity coefficients for Erk (β_E_) and NFAT (β_N_) in the linear models (Fig. 3F). Erk-biased genes include the AP-1 subunits (*Fos*, *Fosb*) and known target genes (*Egr1, Tnf*), as expected (59), whereas genes activated primarily by NFAT included key regulators of exhaustion and effector function (*Pdcd1*, *Lag3, Nr4a3, Ccl3, Ccl4, Ifng*), consistent with known roles of NFAT in driving these gene programs (48). Interestingly, while Erk and NFAT work cooperatively in some modes (A2s, A3s, A3w) and show a positive sensitivity coefficient for combined Erk/NFAT signaling (β_EN_ > 0), they antagonize one another in other modes (A1s, A4s; β_EN_ < 0). Repressed target genes, which include canonical memory and self-renewal genes such as *Tcf7*, *Il7r*, and *Sell*, also showed a range of Erk and NFAT dependencies (R1s-R3s, R1w-R3w, I).

Intriguingly, across all these regulatory modes, we observed a marked negative correlation between the sensitivity to combined Erk/NFAT inputs (β_EN_) and the sensitivities for individual inputs, most prominently for Erk (β_E_) but also for NFAT (β_N_) (Fig. 3H and S3D-H). This pervasive negative correlation suggests a global mechanism generating antagonism between single and combinatorial binding modes. In other systems (60, 61), TFs can be redirected from one set of binding sites to another by a trans-acting binding partner. In T cells, AP-1 and NFAT could redirect each other’s binding from sites where they bind with other partners to those where they bind together. To test whether such a mechanism could explain the inverse relationship between single and combined Erk/NFAT sensitivities, we analyzed candidate mathematical models describing different *cis*-regulatory mechanisms by which AP-1 and NFAT may regulate their target genes (Mathematical Appendix, Section 1). We found that the observed range of sensitivities to single and combined inputs for different target genes (β_E_, β_N_ vs. β_EN_) cannot be readily explained by having composite AP-1:NFAT binding sites alone (Model 1), but most likely reflect binding of these factors at multiple single and composite sites, which could each vary in their transcriptional activity (Model 2). However, having multiple sites alone would give rise to uncorrelated single and combined input sensitivities if these sites act independently from each other to control transcriptional activity. In contrast, when AP-1 and NFAT can redirect one another from single to composite sites, negative relationships between the single (β_E_, β_N_), and combined (β_EN_) input sensitivities arise (Fig. 3H, Model 3), which agrees with experimental data. Importantly, because partner re-direction occurs in *trans* and affects TF availability across all target genes, our model predicts negative correlations would arise across a range of *cis*-regulatory architectures with different input sensitivities, and indeed this is what we observed.

### T cells adopt distinct activation programs in response to variable pMHC affinity and dose

pMHC inputs of different affinity and dose may give rise to distinct gene programs, and do so as a result of decoding of pMHC-dependent Erk and NFAT signaling. To test this, we performed single-cell RNA-sequencing (scRNA-seq) on CD8^+^ T cells cultured with the pMHC conditions we used in our imaging experiments (Fig. 1) using a standard 10X Genomics platform. We analyzed 11,412 cells from five pMHC conditions, as well as a non-stimulated (rest) control. We then performed UMAP dimensionality reduction and clustering of the cells (62, 63) (Fig. 4A) using the 1,089 Erk and NFAT target genes (Fig. 3) identified from bulk RNA-seq.

**Figure 4.**
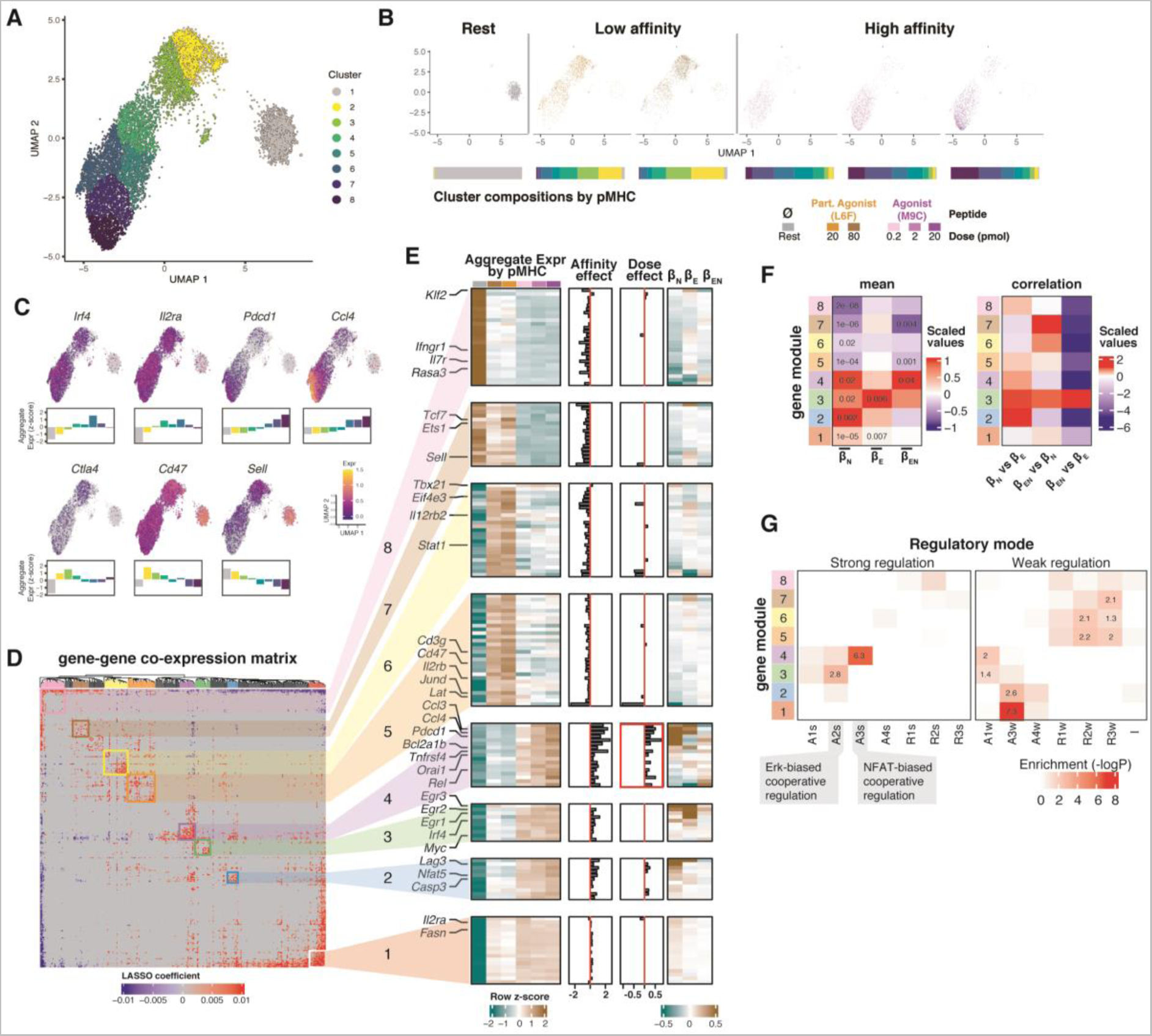
pMHCs of varying affinity and dose elicit distinct T cell functional programs via decoding of Erk and NFAT signaling. (A) UMAP of the 1,089 Erk and NFAT target genes from CD8^+^ T cells stimulated for 30 hrs with five different pMHC conditions. (B) UMAP split by pMHC input. The proportion of cells from each pMHC condition belonging to each cluster in (A) are shown below each UMAP. (C) GOI expression overlaid on UMAP, and their scaled (z-score) aggregate expression by UMAP cluster shown as bar plots. (D) Hierarchical clustering of a single cell gene-gene co-expression matrix of 309 genes against all other genes used in the UMAP analysis. The matrix is colored by the LASSO regression β coefficients for each pair of genes. Each value estimates the magnitude and direction of co-expression between two genes in a single cell. Eight co-expression clusters are identified as gene modules. (E) From left to right: Scaled aggregate expression by pMHC condition for each gene. Normalized effect of pMHC affinity and dose on gene expression determined by differential expression tests in Monocle3. β_N_, β_E_, and β_EN_ values from Fig. 3F for each gene. (F) Summary statistics for β coefficients by gene module. t-test p-values are indicated for mean β coefficients significantly different from zero. (G) Enrichment of regulatory modes identified in Fig 3 within each gene module, with p-values determined by Fisher’s exact test.

Cells rested, or activated with pMHCs of varying affinity and dose localized to different regions of the UMAP and had distinct proportions within each cluster, suggesting the presence of pMHC-specific gene programs (Fig 4B and S4B-C). Amongst activated cells, those exposed to high affinity pMHC (M9C) acquired an effector-like state (Clusters 5-8), with up-regulation of canonical effector genes (*Irf4*, *Il2ra*) and down-regulation of genes associated with memory and lymph node homing (*Tcf7, Sell, Klf2*) (Fig 4C and S4F). Cells exposed to higher doses of this pMHC increased expression of T cell homing chemokines (*Ccl3, Ccl4*); they also expressed higher levels of inhibitory receptors (*Pdcd1*, *Lag3*), suggesting a more rapid progression towards a state of exhaustion (Clusters 7-8). In contrast, cells exposed to low affinity pMHC (L6F) showed less effector differentiation and greater retention of memory programming (Clusters 2-4). Strikingly, these cells enhanced expression of other inhibitory receptors (*Ctla4, Cd47*) as well as *Il2rb* (CD122), a marker of suppressive CD8^+^ T cells (64, 65). These distinct inhibitory receptor profiles suggest multiple suppressive programs depending on the stimulatory regime, and point to a low dose, high affinity pMHC regime for maintaining optimal effector function.

### pMHC affinity- and dose-dependent gene programs reflect decoding of Erk and NFAT signaling

Having identified global differences in the transcriptional states of T cells stimulated with different pMHC inputs, we asked whether these differences arose from the decoding of long-term Erk and NFAT signaling. To do so, we sought to identify genes co-expressed in single cells, as they are likely regulated by the same upstream TFs (63, 66). We used a regression model to compute pairwise gene-gene expression correlations (63), followed by hierarchical clustering on the regression coefficients to identify groups of co-expressed genes we term gene modules (Fig 4D). We found eight gene modules that were either activated or repressed by pMHC stimulation (M1-M6 vs M7-M8) and which varied in both the direction and magnitude of regulation in response to different inputs. Notably, activated modules could be up- or down-regulated with increasing pMHC affinity (M1-4 vs M5-6), or further up-regulated at greater pMHC dose (M4).

To test the role of long-term Erk and NFAT signaling dynamics in generating these pMHC-dependent gene expression patterns, we analyzed the single and combinatorial Erk and NFAT dependencies (β_E_, β_N_, β_EN_) of genes in each module obtained by signaling perturbation measurements (Fig. 4E, right; Fig. 4F and S4E), as well as their enrichment in different Erk/NFAT regulatory modes (Fig. 3F-G; Fig 4G). We found that modules up-regulated with pMHC affinity (M1-M4) showed positive dependencies on Erk/NFAT regulation, whereas those down-regulated with affinity (M5-M8) showed negative dependencies (Fig. 4E-F). These results are consistent with pMHC affinity-dependent differences arising due to stronger Erk and NFAT signaling (Fig. 2E). As increasing pMHC dose further increases NFAT but not Erk signaling (Fig. 2), we predict that modules with strong pMHC dose-dependencies also show greater reliance on NFAT signaling compared to Erk signaling. Indeed, module M4, which showed the greatest pMHC dose sensitivity (Fig. S4D), also showed strong sensitivity to NFAT and the greatest sensitivity to combinatorial NFAT/Erk activity compared with Erk activity (β_N_ >β_E_; β_EN_>β_E_; β_N_>0; β_EN_>0) (Fig. 4F). Furthermore, this module was enriched for genes exhibiting an NFAT-biased mode for cooperative regulation (Fig. 3, mode “A3s”; Fig. 4G), thus identifying composite NFAT:AP-1 regulation as a regulatory mode for sensing pMHC dose. Similar NFAT-dominant regulation was also observed for the moderately dose-dependent module M2 (β_N_ >β_E_; β_N_>0), though, interestingly, this module showed considerably weaker sensitivity to combined Erk/NFAT regulation (β_EN_). In contrast, module M3, which was strongly up-regulated with increasing affinity but showed no dose-dependency, showed the greatest sensitivity to Erk signaling (β_E_), and was also enriched for genes exhibiting an Erk-biased cooperative mode of regulation (mode “A2s”). This finding aligns with the pMHC affinity dependence and dose independence of this module, given that late Erk activity is the largest signaling difference between high and low affinity pMHC (Fig. 1-2). Additionally, some modules showed weaker sensitivities to Erk and/or NFAT signaling, yet they showed distinctive dependencies on pMHC inputs (M1, M5, M8), suggesting the presence of additional *trans*-factors responsible for decoding pMHC inputs. Taken together, these results indicate that pMHC affinity- and dose-specific T cell states arise, at least in part, from decoding of long-term signaling dynamics by Erk and NFAT target genes.

### Distinct *cis*-regulatory mechanisms underlie the decoding of pMHC-dependent Erk and NFAT signaling

To gain insights into the mechanisms integrating Erk and NFAT signaling inputs to generate pMHC-specific expression patterns, we analyzed mathematical models of candidate *cis*-regulatory mechanisms for two well-studied T cell activation genes, *Pdcd1* (encoding PD-1) and *Il2ra* (encoding CD25), to account for experimentally measured Erk and NFAT signaling responses and gene expression patterns. These genes were chosen for their distinct pMHC affinity and dose dependencies, as well as their inclusion in distinct modules (M4 for *Pdcd1*, M1 for *Il2ra,*), allowing us to explore a range of *cis-* regulatory mechanisms for decoding signaling dynamics (Fig 5A). This approach utilizes quantitative measurements of both processes to identify and constrain models of *cis*-regulatory element function, allowing us to determine whether the modeled mechanisms can explain experimental observations, and to determine the plausible parameter regimes under which these mechanisms operate. We used flow cytometry to measure PD-1 and CD25 levels in T cells activated by M9C and L6F at different doses, as above (Fig 5C-D and S5). We also perturbed cells with Erk and NFAT inhibitors after 9 hours of stimulation (“Late”). However, because some *cis*-regulatory elements can control activation timing without affecting maintenance of gene expression (61, 67–69), as observed for *Il2ra* (67), we also perturbed signaling at the onset of stimulation (“Early”), as Erk and NFAT may play distinct roles in controlling activation versus maintenance of gene expression. Based on these measurements, we developed and fit these data to dynamical models that describe the *cis*-regulatory control of these two genes (see Mathematical Appendix).

**Figure 5.**
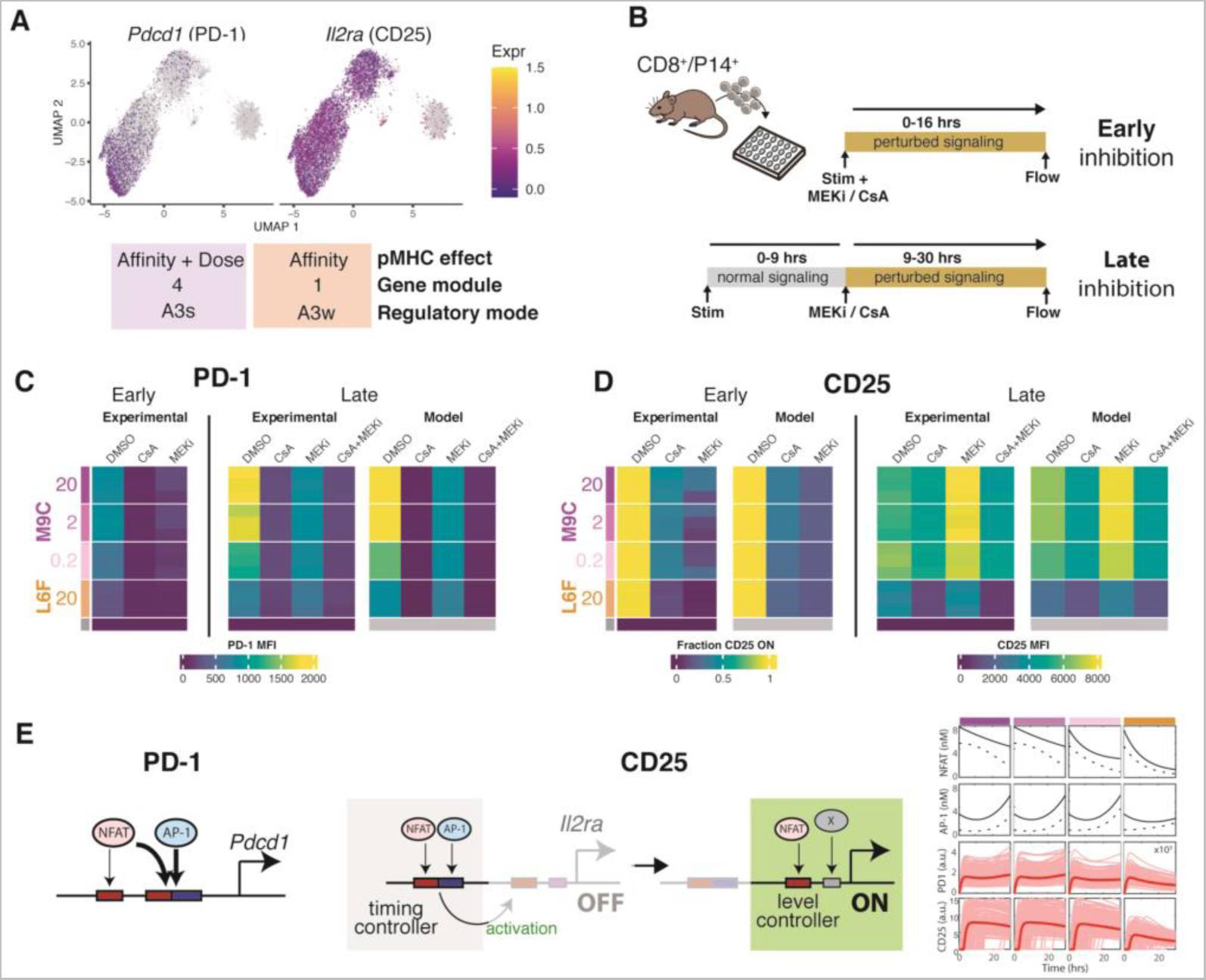
Temporal and combinatorial *cis*-regulatory mechanisms work together to decode Erk and NFAT signaling. (A) scRNA-seq UMAPs overlaid with *Pdcd1* and *Il2ra* expression. (B) CD8^+^ T cells were stimulated with various pMHC conditions, treated with MEKi and/or CsA immediately (early) or 9 hrs (late) after stimulation, then assayed for PD-1 and CD25 expression after 16 (early) or 30 (late) hrs. (C-D) Expression of PD-1 and CD25 in T cells following variable stimulation and signaling perturbation shown in (Β). Experimental results are compared with our model predictions. (E) Proposed NFAT and AP-1 *cis*-regulatory mechanisms controlling PD-1 (*Pdcd1*) and CD25 (*Il2ra*) expression. Mathematical simulations for each pMHC input are shown for each gene, depicting the levels of active NFAT and AP-1 transcription factors, and the resulting expression of PD-1 and CD25.

This analysis revealed an array of regulatory mechanisms that collaborate in *cis-* and *trans*- to enable input-specific expression of these two genes downstream of Erk and NFAT signaling. PD-1 expression is predicted to depend on the concerted action of two elements, both of which are required for the observed dose, affinity and inhibitor dependencies in its expression: (1) an element that binds NFAT without AP-1, and (2) a composite NFAT:AP-1 binding site that acts independently (Fig 5E). This predicted bipartite architecture contrasts with an alternative where partnerless NFAT may play a dominant role in *Pdcd1* regulation (48), and highlights how target genes may utilize multiple elements that bind TFs in different combinations to construct input-specific responses. CD25 expression is also predicted to require two separate elements, but that have distinct regulatory functions: (1) a timing element consisting of an NFAT:AP-1 composite binding site, and (2) an element controlling expression levels that receives input from NFAT and an undetermined factor X (Fig 5E). As the timing element is required for initiation but not maintenance of expression, this system enables hysteresis in its regulation, allowing target genes to activate in response to prior signaling exposure. Interestingly, after activation, CD25 expression is regulated not only by a third, unidentified factor, but also by NFAT whose effects are revealed only upon attenuation of Erk signaling and presumably occur by NFAT redirection by AP-1 at other gene loci (Fig 5D and S5B). These results highlight how *trans*- mechanisms for TF partner redirection could generate additional dependencies for decoding combinatorial signaling states generated by diverse signaling inputs.

## DISCUSSION

Using an *in vitro* platform to follow individual T cells responding to varying pMHC inputs, we find that T cells independently encode pMHC affinity and dose information using the long-term (9+ hrs) dynamics of these two pathways.. Through transcriptomic profiling, we find that Erk and NFAT signaling dynamics are decoded to generate pMHC affinity and dose specific gene programs, with multiple mechanisms by which these inputs are integrated at the *cis*-regulatory level. This work reveals a previously unappreciated role for long-timescale signaling in antigen encoding, and highlights the combinatorial effects of multiple signaling pathways in encoding distinct signaling input features.

Our findings indicate that T cells initially activate uniformly, and elaborate pMHC-specific signaling and gene regulatory responses only after prolonged contact with various pMHC inputs. This shared molecular program for early activation, also observed in other studies (25, 26), may enable T cells to prepare for a potential response before fully determining the nature of the threat. Such early response initiation may allow for rapid responses if a threat is indeed present, which may be critical as T cells reach the point of first cell division only ∼30 hrs after pMHC encounter (70). The lag in pMHC dose and affinity-dependent responses may be due to time-delayed negative feedback on signaling resulting in reduced sensitivity to pMHC inputs, such that only high affinity pMHC can overcome this feedback. Indeed, T cell signaling is subject to multiple layers of negative feedback (6, 71, 72). Some of these negative feedback loops, particularly those involving transcription of checkpoint inhibitory molecules (73, 74), may act on extended time-scales similar to those we observed. Alternatively, the pMHC-specific response lag may reflect a need for T cells to integrate pMHC information over extended durations for accurate input discrimination (72, 75, 76). In this picture, late divergent Erk and NFAT signaling levels would result from prior pMHC exposure along with immediate signaling inputs, potentially through cytokine feedback loops that could provide additional inputs into these pathways (22, 72).

The multiple mechanisms by which gene regulatory elements can decode pMHC-dependent Erk and NFAT signals enables T cells to elaborate distinct functional programs in response to different pMHC inputs. Our transcriptomic studies of Erk and NFAT regulation, together with further in-depth analysis of *cis*-regulatory systems of specific genes (*Pdcd1* and *Il2ra*), indicate that target genes can be controlled through multiple *cis*-regulatory elements where Erk and NFAT can either act in combination, or singly; each of which may have different transcriptional outputs. Furthermore, Erk and NFAT can be readily redirected on a global level by the abundance of their partners, also observed in other contexts (60, 77), and may shift the usage of these *cis-*regulatory elements depending on the cell’s signaling state. Finally, Erk and NFAT signaling activity can modulate gene activation timing but not maintenance of expression, possibly through enhancer elements that control chromatin states (28, 61, 67, 78), a finding that provides a mechanism by which cells can temporally-integrate cumulative Erk and NFAT signaling dynamics to generate stable pMHC-dependent changes in gene programming (68).

An ability to independently perceive pMHC affinity and dose may be critical for allowing T cells to selectively counter pathogens while suppressing responses to self that could lead to autoimmunity. While T cells with high affinity for self-pMHCs are eliminated during negative selection, those retained in the periphery can have affinities for self-pMHC that are very close to the sharp threshold for negative selection(3, 79, 80). Though these T cells with lower-affinity TCRs are able to fully activate upon antigen encounter (25, 26), they are far less efficient in mediating autoimmune destruction of healthy tissues compared to those with high-affinity TCRs (81), but may rely on a suppressive program, nonetheless, to prevent spurious effector responses. Our transcriptomic profiling showed that T cells activated by low-affinity pMHCs maintain a distinct functional program with attenuated effector programming (*Irf4* and others) and up-regulation of the immune checkpoint and suppressor function genes (i.e. *Ctla4*, *Cd47*, *Il2rb*). In contrast, T cells activated by high doses of high-affinity pMHCs did not activate these suppressive genes, but instead upregulated distinct immune checkpoint genes (e.g. *Pdcd1*, *Lag3*), possibly reflecting a distinct inhibitory program for dampening hyperactive cytotoxic activities during severe pathology. Future studies should investigate these pMHC-dependent regulatory and functional states, particularly in contexts where T cell antigen recognition can result in initiation of autoimmune pathologies.

## MATERIALS & METHODS

### Data Availability

Further information and requests for resources and reagents, including mouse lines generated in this study should be directed to and will be fulfilled by the corresponding author, Hao Yuan Kueh.

### Mice

The dual-pathway reporter mouse strain (*Rosa26^LSL-DPR/LSL-DPR^*) was generated on a B6 background by Biocytogen Corp. using a *Rosa26* targeting vector (82) containing a reporter cassette encoding Erk-KTR (53) fused to mTagBFP2, histone 2B fused to iRFP, and residues 1-399 of mouse NFATc2 fused to mRuby3 (51). *Rosa26^LSL-DPR/LSL-DPR^* mice were crossed to LCMV-specific transgenic TCR (P14) mice (83) (Jackson Labs, Strain # 004694) and to Cre-ERT2 mice (Jackson Labs, Strain # 008463), generating *Rosa26^LSL-DPR/Cre-ERT2^/TcrLCMV* offspring. 8 week old male and female mice were injected with four doses of tamoxifen over four consecutive days (84) and used for imaging experiments 4-12 weeks later. 8-16 week old *Rosa26^LSL-DPR/LSL-DPR^/TcrLCMV* male and female mice were used for non-imaging experiments (Fig. S1A). All mice were used in accordance with Institutional Animal Care and Use Committee guidelines for the University of Washington.

### pMHC-coated plate*s*

Glass-bottom (MatTek Cat. no. PBK96G-1.5-5-F) or TC plates were prepared as previously described (55) with minor modifications. Plates were coated with 2 ug Retronectin (TaKaRa Cat. no. T110A), 1 ug anti-mouse CD11a (LFA1), and 2 ug NeutrAvidin (Thermo Fisher Scientific, Cat # 31000) in 100 ul PBS per well and incubated overnight at 4°C. Retronectin and anti-LFA1 reduce T cell motility and enable long-term tracking (85). Plates were blocked with 1% BSA for 1 hr at 37°C, then incubated with pMHC for 90 minutes at 4°C. Biotinylated monomeric H-2D^b^ MHC loaded with KAVYNFATM (M9C) or KAVYNLATC (L6F) (57) were provided by the NIH tetramer facility and diluted in cold PBS. The molar quantities of pMHC indicated in the text and figures corresponds to the total amount of pMHC incubated in each well in 100 ul PBS (i.e. 100 ul of 10 ug/ml pMHC solution ∼ 20 pmol pMHC).

### T cell stimulation

CD8^+^ T cells were isolated from RBC-depleted splenocytes using the Miltenyi CD8^+^ negative selection kit (Miltenyi Biotec, Cat # 130-104-075). Reporter-expressing CD8^+^ T cells were sorted using a BD Aria III. For dose-response titration experiments, a monoclonal P14 CD8^+^ Jurkat T cell line was used, and was generated by transducing *Tcrb*-KO Jurkat T cells (a gift from Phil Greenberg) with lentiviruses containing the P14 TCRα and TCRβ chains, and mouse CD8α and CD8β1 chains. Cells were cultured in AIM-V media (Thermo Fisher Scientific, Cat # 12055091) supplemented with 50 uM BME. Rested cells for scRNA-seq and flow cytometry experiments were cultured in media supplemented with 10% heat-inactivated FBS, 5 ng/ml hIL-7 (Peprotech Cat # 200-07), and 50 ng/ml mIL-15 (Peprotech Cat # 210-15). Seeded cells were spun at 80g for 2 min and incubated at 37°C with 5% CO_2_.

### Live-cell imaging

Time lapse images were acquired using a 63X (0.75 NA) glycerol objective on a Leica DMi8 equipped with hardware autofocus and a 37°C, 5% CO2 incubation chamber. Images were taken with a Photometrics Prime 95B camera using automated software (MetaMorph, Molecular Devices) to acquire DIC images every 20 min and fluorescence images every 60 min. Fluorescence images were acquired in three channels: BFP (405 Ex; 440/40 Em), RFP (561 Excitation; 600/50 Em), and near-IR (640 Ex, 700/75 Em) using an LED illuminator (SpectraX, Lumencor).

### Flow cytometry

Antibodies used for flow cytometry can be found in the SI. Cells were stained in Fc blocking solution (2.4G2 supernatant) containing: 1 ul anti-CD69, 0.5 ul anti-CD25, or 1 ul anti-CD279 (PD-1) per sample and analyzed on an Attune Nxt cytometer (Thermo Fisher Scientific).

### Bulk RNA sequencing

450k CD8+ T cells per condition were stimulated with 2 pmol M9C. After 9 hours, cells were treated with either 1 uM MEK inhibitor (Trametinib, Selleckchem Cat. no. S2673), 1 uM Cyclosporin A (Selleckchem Cat. no. 52286), 1 uM of both, or DMSO (0.1%), and cultured for an additional 21 hours. Cells were then washed 1x in ice-cold PBS, resuspended in 350 ul Trizol (Invitrogen Cat # 15596018) and vortexed for 30 sec to lyse the cells. RNA was isolated with the RNeasy Micro kit (Qiagen Cat # 74004). Paired-end library preparation and transcriptome sequencing was conducted by Novogene Co., LTD (Sacramento, CA, USA).

### Single-cell RNA sequencing

Following 30 hrs of stimulation, cells were stained with one of the six hashtag (HTO) antibodies (1 ug HTO per sample) for 30 min at 4°C. 22k live cells (4k from each pMHC condition + 2k rested cells) were flow sorted using propidium iodide (Alfa Aesar, Cat # J66584). Single cell library preparation was done with the 10x Genomics platform, loading all 22k cells into a single lane (86), and following the manufacturer’s protocol (CG000205, Rev D, Single Cell 3’ v3.1 with feature barcoding, 10x Genomics). Libraries were diluted to 2nM using Agilent Tapestation D1000 HS and loaded onto two NextSeq 2000 P2 100-cycle kits using the following conditions: read 1: 28 cycles; index 1: 8 cycles; read 2: 92 cycles. Data was converted to fastq using bcl2fastq. Cellranger (version 6.0.1, 10x Genomics) was used for transcriptome-(337M reads) and HTO-(161M reads) level unique molecular identifier (UMI) and cell demultiplexing, and for aligning the transcriptome to the GRCm38/mm10 reference genome. Doublets and cells with > 7.5% mitochondrial reads were removed.

### Image processing

Image pre-processing, cell segmentation, and tracking was performed with a custom MATLAB (Mathworks) pipeline as described previously (69) (Fig. S1B). Nuclear masks were generated by segmenting H2B-iRFP images. Cytoplasmic masks were generated by first creating a weighted average image of all three fluorescent images, segmenting the resulting image to yield a total cell mask, then subtracting the nuclear mask. Cells are then linked from frame to frame to generate single-cell traces of signaling activity, which were manually validated. We calculated the median nuclear fluorescence and the 0.75 quantile of cytoplasmic fluorescence, since we observed uneven distribution of fluorescence in the cytoplasm, and determined pathway activities as follows:

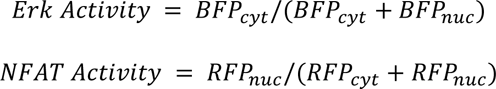

### Computational analysis of Erk and NFAT signaling

#### Single cell dual-pathway clustering

Bivariate 14-dimensional vectors (14 timepoints for each pathway) were scaled from zero to one and clustered with the *dtwclust* package (v 5.5.11) (87) in R using the global alignment kernel distance (88), which allows for calculating distances between multivariate single cell responses. Parameters used with the *tsclust* function were: k = 8, type = “partitional”, distance = “gak”. The use of 8 partitions was empirically determined.

#### Quantification of dynamic features

Occurrence of a second signaling pulse was determined using two measures: (1) a 3 hr sliding window of time-integrated areas to further smooth the single cell activity traces (A_smooth_), and (2) the derivative of the smoothed trace (dAdt). A pulse was detected in each pathway if there were at least 2 consecutive hours with dAdt > 0 and A_smooth_ > (0.95 x Ā_smooth_). Ā_smooth_ was calculated as the mean smoothed activity for each pathway from all analyzed cells. Pulse time (t_pulse_) was defined as the latest positive dAdt within the pulse window, and pulse amplitude (A_pulse_) was defined as the smoothed activity value at t_pulse_,and then scaled (z-score). Early and late integrated activities for each pathway were calculated using the activity traces prior to the pulse detection smoothing, and computed using the 2-4 hr and 8-15 hr activities, respectively.

#### Machine learning classification models

We used a 5-fold nested cross-validation training and validation workflow (89) whereby for each fold, a random forest classification model with 500 decision trees was trained and subsequently validated on the hold out partition using the *randomForest* package (v 4.7-1.1) in R.

### Computational analysis of RNA sequencing data

#### Bulk RNA-seq DEG analysis

Raw FASTQ files were aligned to the GRCm38/mm10 reference genome using Kallisto (v0.46.1) (90). Transcripts with < 5 counts were excluded using the *sleuth* package (v 0.30.0) (91) in R. Differential expression analysis was performed with *sleuth* following the Pachter lab walkthrough (https://pachterlab.github.io/sleuth_walkthroughs/boj/analysis.html), setting the “MEKi+CsA” treatment as the control. We classified DEGs based on a q-value threshold of 0.001. To enable comparison with scRNAseq, we took the intersection of this DEG list (1,241 genes) and the list of expressed genes from the scRNA-seq dataset after quality control filtering to yield 1,089 DEGs for our analyses. We performed Wald tests on each β coefficient (MEKi, CsA, DMSO) from the “full” model to determine Erk- and NFAT-regulated genes using a 0.001 q-value cutoff.

#### Multiple linear regression models

We fit a linear model to each of the 1,089 DEGs using the *lm* function within the *stats* package (v 4.1.2) in R with the following formula: 𝑦 ∼ 𝑥_𝐸_ ∗ 𝑥_𝑁_, where ŷ_𝑔𝑒𝑛𝑒_ is the normalized expression upon inhibitor treatment, and 𝑥_𝑁_, 𝑥_𝐸_ are the NFAT and Erk integrated activities upon treatment (Fig. S3C).

#### scRNA-seq UMAP clustering

Quality control, pre-processing, dimensionality reduction, and UMAP visualization were done with *Monocle3* (92) (v 1.2.9) in R following the Trapnell Lab walkthrough (https://cole-trapnell-lab.github.io/monocle3/docs/clustering/). The scRNA-seq cell dataset was subset on the bulkRNA-seq DEGs expressed in at least 100 cells, generating a cell dataset contained 1,089 genes and 11,956 cells, with a median UMI/cell count of 5,972. Cells were partitioned into 8 clusters using the *cluster_cells* function. These 8 clusters contained 95% of the cells, with additional outlier clusters containing the remaining 5% (544 cells) which were removed for all further analyses.

#### Single cell gene-gene co-expression analysis

We performed LASSO (least absolute shrinkage and selection operator) regression using the *glmnet* package (v 4.1-4) in R following a previously developed analysis method (63) on the 810 genes expressed in at least 5% of cells.

### Mathematical modeling

Ordinary differential equations simulations of gene regulation models for *Pdcd1* and *Il2ra* were performed using MATLAB. Parameter values for these models were determined by the fitting gene expression values obtained by modeling to experimental measurements of mean fluorescence intensities (*Pdcd1* MFI, *Il2ra* late, Fig. 6C, D) or gene activation percentages (*Il2ra* early, Fig 6D). The sum-squared error of this fit was then minimized using the Nelder-Mead simplex method. In these models, cell-to-cell variability in gene expression was modeled using a cell-extrinsic distribution, via log-normally distributed differences in maximal transcriptional activity. In our model, activation times for *Il2ra* were exponentially distributed. For simulations, these random activation times were determined through random number generation, followed by the distribution of likelihoods.

## Acknowledgments

We thank members of the Kueh and Fowler labs for discussion and feedback, Tony Cooke (previously with Leica Microsystems) for microscope setup and support, Philip Greenberg for providing *Tcrb*-KO Jurkat cells, and the NIH Tetramer Core Facility for providing pMHC monomers. This study was funded by an NIH/NIBIB Trailblazer Award (R21EB027327, H.Y.K.), an NIH/NHGRI Program Project Grant (RM1HG010461, D.F.), a NSF Graduate Research Fellowship (M.J.W.), and startup funds from the Bioengineering Department at the University of Washington (H.Y.K.).

