## Supplementary Information for "Antigen perception in T cells by long-term Erk and NFAT signaling dynamics"

**Supporting Information**

**This PDF file includes:**

Figures S1-5

Legends for Supplementary Videos 1-5

Legends for Supplementary Tables 1-2

Supplementary Methods

Mathematical Appendix

Supplementary References

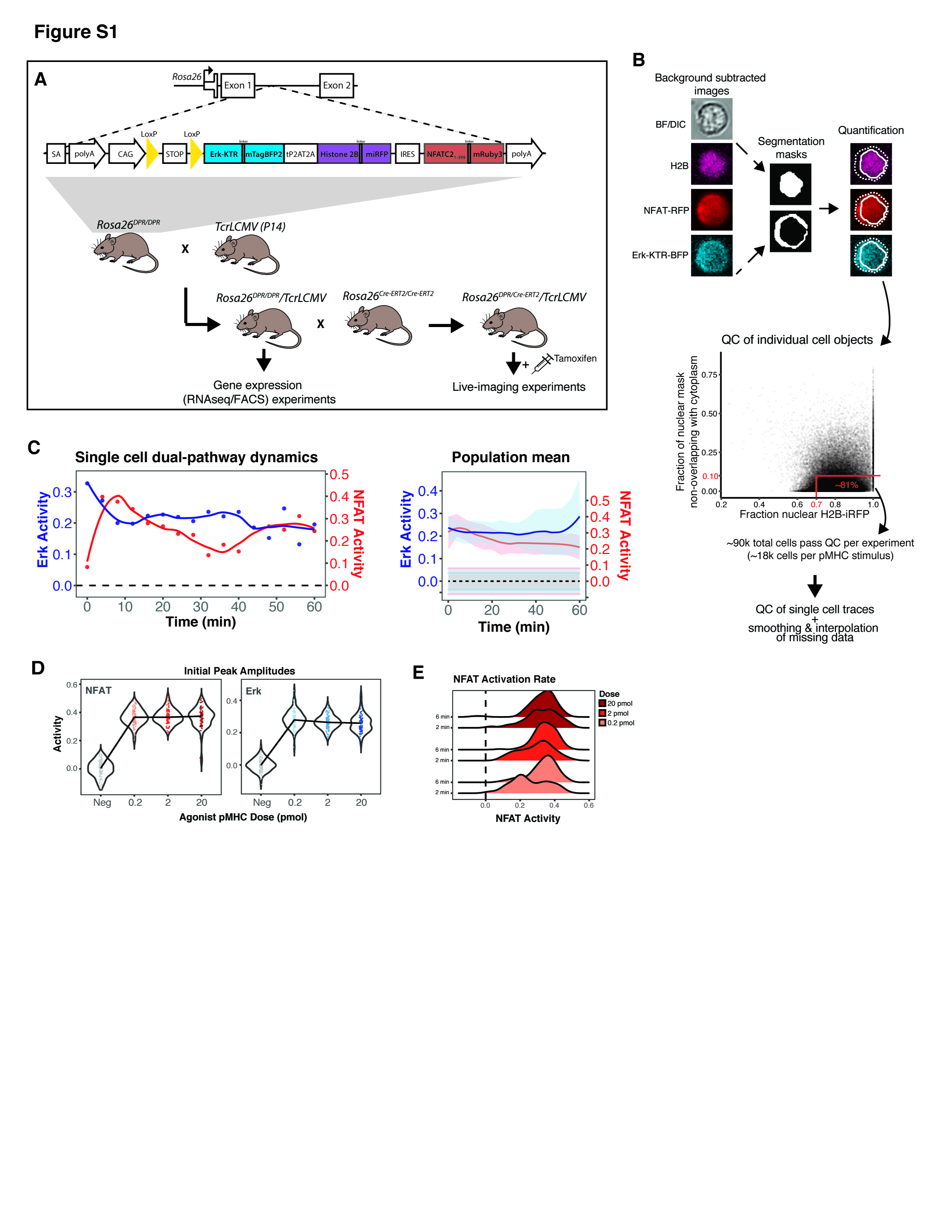

**Figure S1. Dual-pathway reporter mouse and automated image analysis pipeline enables quantification of Erk and NFAT signaling dynamics in single T cells.**

**(A)** Schematic of the reporter transgene insertion into the mouse Rosa26 locus. Homologous recombination was done by Biocytogen Corp. using their CRISPR-based EGE^TM^ method. Breeding strategy used to generate all mice used in the study, indicating which strains were used for which experiments.

**(B)** Overview of MATLAB image processing and quantification workflow. Quality control and single cell analyses were done in R.

(C) Quantification of Erk and NFAT activity for the cell shown in (Fig 1B). Population mean +/- SD from 107 stimulated cells is shown on the right. Pathway activities are baseline subtracted using the mean pathway activities from negative control cells and shown as dashed lines with SD in gray shading.

**(D)** Reporter T cells stimulated with a 100-fold dose range of agonist pMHC (M9C) show invariant initial peak amplitudes for both Erk and NFAT signaling responses, as determined by the maximum activity in the first 12 minutes (4 frames) of stimulation.

**(E)** Distribution of NFAT activities between the first (~2 min) and second (~6 min) frames are shown for the three doses indicated. The fraction of cells that activate NFAT within 2 minutes of TCR stimulation is graded and increases with increased pMHC dose. By 6 minutes, all doses produce similar NFAT activation.

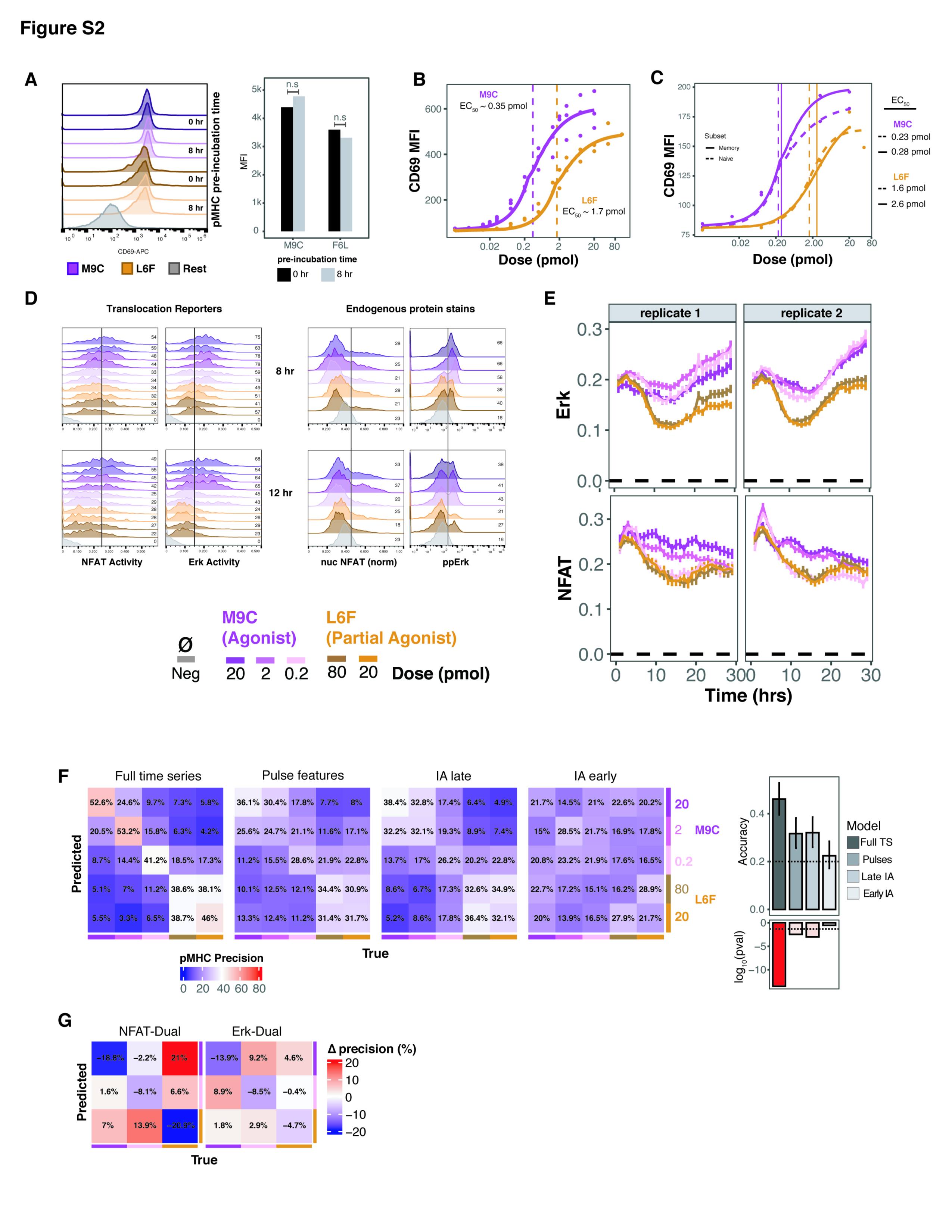

**Figure S2. Validation of Erk and NFAT signaling assay and independent encoding of pMHC affinity and dose information by long term Erk and NFAT signaling.**

**(A)** Both APLs (M9C and F6L) used in this study remain active (i.e properly folded) for at least 8 hrs at 37^0^C. T cells were assayed for CD69 expression 4 hrs after stimulation on pMHC-coated plates that were pre-incubated with 20 pmol of each APL for 8 hrs at 37^o^C, or coated immediately prior to cell seeding. No difference in CD69 MFI was observed between pre-incubation times for each APL, as determined by t-test.

**(B)** Raw CD69 MFI dose-response curves for P14 Jurkat T cells for both APLs used. EC_50_s are indicated in dashed lines. Flow analysis is done after 4 hrs of stimulation.

**(C)** CD69 dose-response curves, as in B, for naive (CD62L^+^/CD44^-^) and memory (CD62L^+^/CD44^+^) murine P14 CD8^+^ T cells. EC_50_s are indicated in dashed and solid vertical lines.

**(D)** Comparison of Erk and NFAT activity measurements between live translocation reporters and endogenous NFAT1 and ppErk staining. For endogenous protein staining, T cells were stimulated identically to cells stimulated for live imaging and fixed, permeabilized, and stained for NFAT1 or ppErk at 8 and 12 hrs post stimulation. Non-stimulated cells were used as a negative control for endogenous protein staining. Nuclear NFAT1 was measured with fluorescent microscopy and normalized to total NFAT1 fluorescence. ppErk intensity was measured with flow cytometry. Vertical lines are shown for reference, with the fraction of cells to the right of the line indicated.

**(E)** Our live-cell imaging and automated image analysis pipeline is robust and reproducible based on results from two independent experiments performed several weeks apart with separate litters of mice. Plots show mean +/- 99% CI Erk and NFAT activity over 30 hrs of stimulation from ~90k total cells per experiment stimulated with five different pMHC conditions (same as in Fig 2C).

**(F)** Random forest model precision in predicting pMHC input considering all five pMHC conditions assayed (compared to only three of the five in Fig 2), and model accuracies and p-values testing for greater accuracy compared to that by random chance (0.2 accuracy).

**(G)** The precision matrix resulting from the “Full time series”-trained random forest classification model was subtracted from those of models trained on either the NFAT (“NFAT-Dual”) or Erk (“Erk-Dual”) time series responses. The resulting Δprecision matrices indicate the effect of removing information from the Erk (“NFAT-Dual”) or NFAT (“Erk-Dual”) pathways.

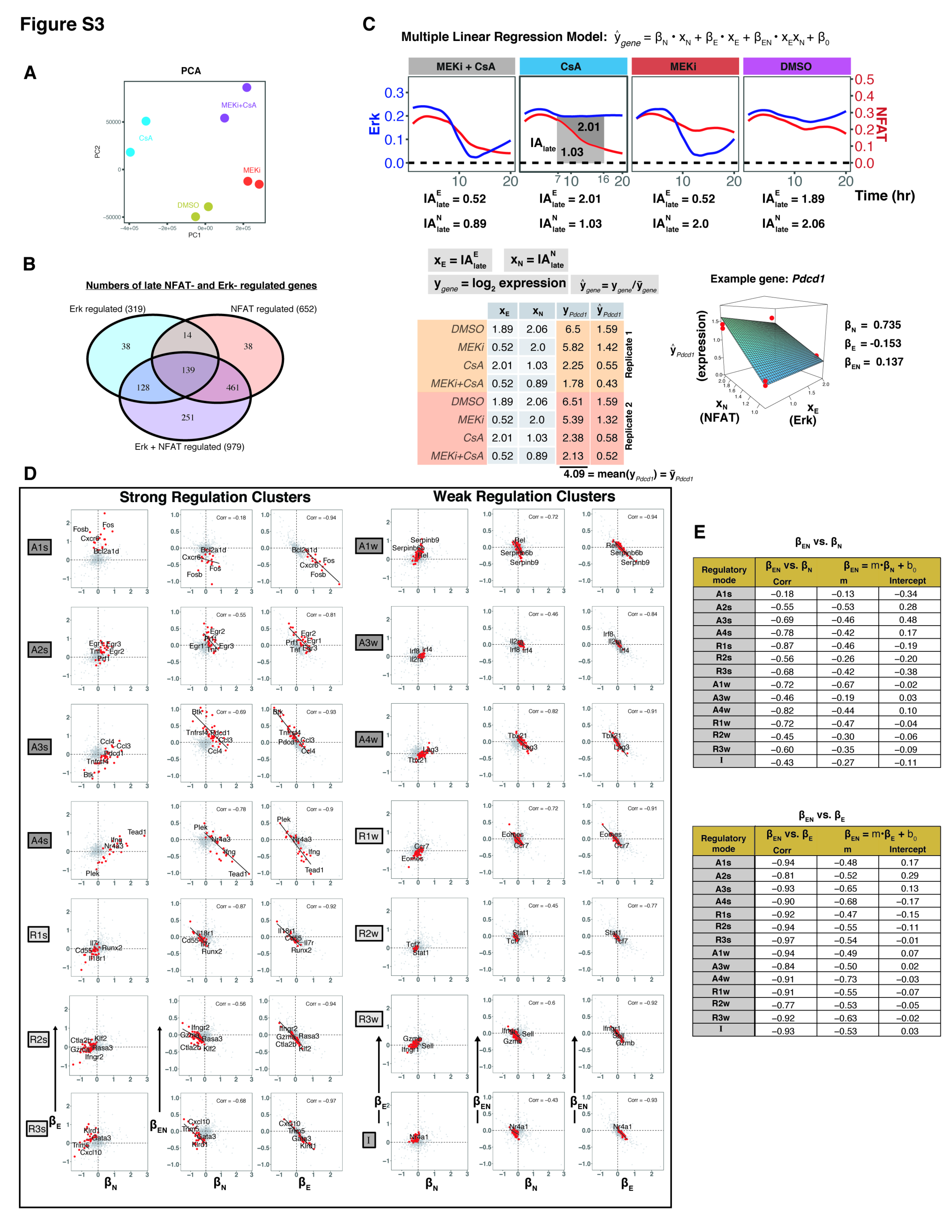

**Figure S3. Characterization of distinct modes of Erk and NFAT decoding.**

**(A)** Principal component analysis performed on bulk RNAseq data comparing T cells treated with Erk (MEKi) and NFAT (CsA) inhibitors from 9-30 hrs of stimulation.

**(B)** Numbers of differentially expressed genes that are regulated by NFAT, Erk, or both. Gene lists can be found in the supplemental tables.

**(C)** Overview of the method for the multiple linear regression models presented in Fig 4F. The integrated activity from 7-16 hrs was calculated from the mean traces of Erk and NFAT activity from cells treated with DMSO, MEKi, CsA, or MEKi+CsA. These values served as predictor variables for modeling the expression of each gene under the same inhibitor conditions, which was determined by bulk RNA seq. Calculation of β coefficients for *Pdcd1* is shown as an example.

**(D)** Pairwise correlations between β coefficients for each regulatory mode is plotted separately as scatter plots with genes comprising each regulatory mode shown in red (all other DEGs shown in gray). Linear regressions and Pearson correlation coefficients are shown for β_EN_ vs. β_N_ and β_E_.

**(E)** Table listing Pearson correlation coefficients and linear regression coefficients for β_EN_ vs. β_N_ and β_E_ for all regulatory modes identified from the bulk RNA seq hierarchical clustering analysis.

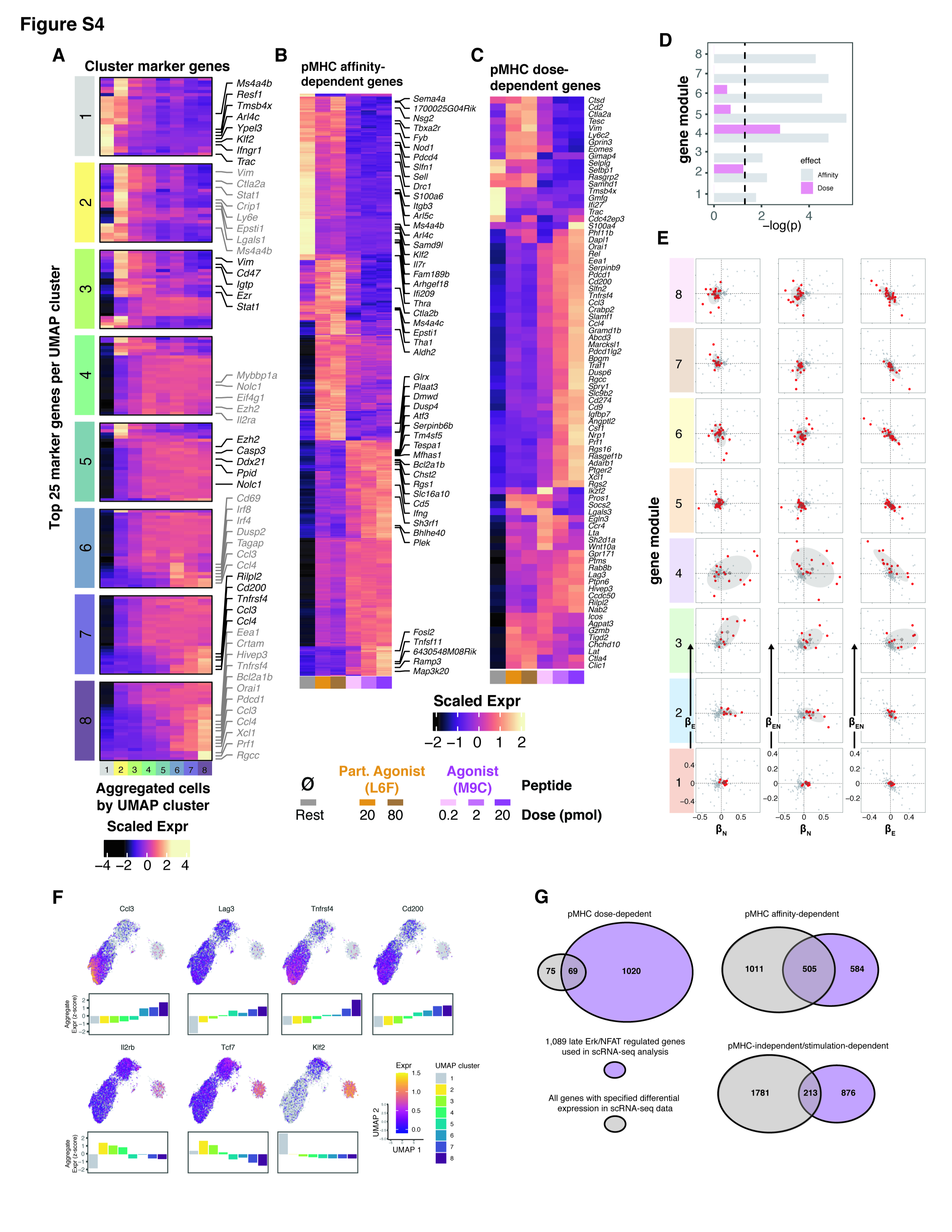

**Figure S4. pMHC-dependent gene expression responses.**

**(A)** Heatmap of the top 25 marker genes for each UMAP cluster from Fig5A. The top 5 marker genes for each cluster are annotated, along with genes that are unique to that cluster. Expression of each gene is aggregated by cluster, and scaled as a z-score by row. Coloring of annotations serves only to aid in distinguishing annotated genes from adjacent clusters. See supplementary tables for full gene list.

**(B)** Heatmap of 591 pMHC affinity-dependent Erk and NFAT target genes identified from DEG tests using Monocle3. Expression of each gene is aggregated by pMHC condition, and scaled as a z-score by row (color legend is shared with C). The top 50 DEGs based on effect size are annotated. See supplementary tables for full gene list.

**(C)** Heatmap of 82 pMHC dose-dependent Ek and NFAT target genes identified from DEG tests using Monocle3. Expression of each gene is aggregated by pMHC condition, and scaled as a z-score by row.

**(D)** Wilcoxon signed rank tests comparing the affinity and dose dependencies of each gene module in Fig. 5D-E. All modules show affinity-dependence based on a p-value threshold of 0.05 (dashed line), while only cluster 4 is dose-dependent.

**(E)** Pairwise correlations between each β coefficient for all gene modules are plotted separately as scatter plots with genes comprising each module shown in red, and all other genes in the co-expression matrix from Fig 5D shown in gray. The 1 SD (67%) confidence high density region is highlighted with a gray ellipse.

**(F)** Expression of select genes of interest overlaid on UMAP, and their scaled (z-score) aggregate expression by UMAP cluster shown as bar plots.

**(G)** Venn diagrams visualizing the overlap of Erk and NFAT target genes (1,089 DEGs identified from bulk RNA-seq) vs all genes identified in the scRNA-seq dataset as being pMHC affinity or dose dependent, or pMHC independent. See supplementary tables for full gene lists.

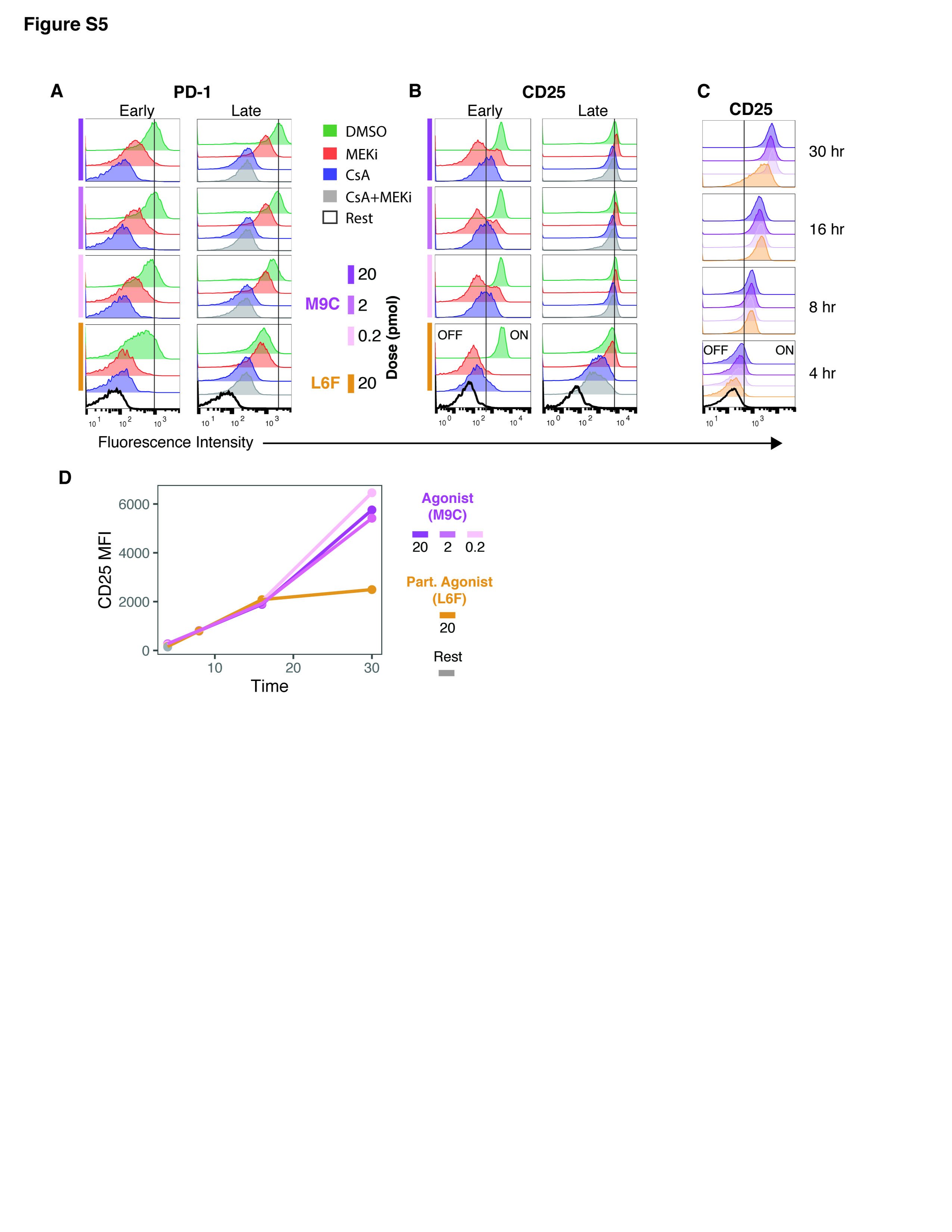

**Figure S5. Regulation of PD-1 and CD25 by Erk and NFAT signaling.**

**(A-B)** Flow cytometry plots of PD-1 and CD25 expression in response to different pMHC inputs, and treated with MEKi and CsA at early or late timepoints. See Fig 6B for experimental design. pMHC condition is indicated with the colored bars on the left. Black lines are drawn for reference, with the line for early CD25 response demarcating CD25 ON vs OFF cells.

**(C)** Time course of CD25 expression in response to different pMHC inputs. Histograms are colored by pMHC condition. Black line shows CD25 ON vs OFF gate from B (“Early”).

**(D)** MFI of CD25 expression over time from the data in (C).

**Supplementary Videos 1-5 (separate files).** **Live-cell microscopy of T cell stimulation.** Representative time-lapse movies of individual well positions corresponding to each of the five pMHC conditions assayed in this study. Shown to the right of the movies for each condition is the mean +/- 99% confidence interval Erk and NFAT signaling responses for all cells analyzed from the specific well position shown in the movie. Nuclear and cytoplasmic segmentations are shown in dashed and solid lines, respectively. Scale bar = 5 um.

**Supplementary Table 1 (separate file).** **DEGs and linear models from bulk RNA-seq data**. The column “Detected in scRNAseq data” indicates if the gene was part of the 1,089 Erk- and NFAT-regulated genes used in our analyses. “gene fits” are the multiple linear regression model parameters for each of the 1,089 Erk and NFAT regulated genes. Provided are the 8 fitted, normalized log_2_ expression counts from two replicates of the four treatment conditions for each gene, the β coefficients (Erk, NFAT, ErkNFAT) along with their p values and standard errors, the model intercept (B0), R^2^, and residual standard error (RSE) values.

**Supplementary Table 2 (separate file). DEGs from scRNA-seq data analysis**. Marker genes for each of the eight UMAP clusters, determined with the *top_markers* function in *Monocle3*. Ordered list of genes comprising the gene-gene correlation matrix following LASSO regression analysis. Gene inclusion in a gene module is indicated. DEGs comparing pMHC stimulation to rest. pMHC affinity-dependent DEGs comparing 2 and 20 pmol M9C stimulation to 20 and 80 pmol L6F stimulation. pMHC dose-dependent DEGs comparing all three doses of M9C stimulation. pMHC affinity-dependent DEGs regulated by Erk/NFAT determined by the intersection of the full affinity-dependent gene list with the 1,089 Erk/NFAT-regulated genes. pMHC dose-dependent DEGs regulated by Erk/NFAT determined by the intersection of the full dose-dependent gene list with the 1,089 Erk/NFAT-regulated genes.

**SUPPLEMENTARY METHODS**

**ppErk Staining**

For ppErk staining, cells were fixed in 2% formaldehyde (Fisher Scientific Cat # 50-980-487) in PBS for 20 min at RT. Cells were washed 1x and stored at 4^o^C until all time points were harvested, then permeabilized with ice-cold 90% MeOH at 4^o^C overnight (36). Next, cells were washed 1x, blocked for 15 min at 4^o^C with Fc blocking solution. 0.1 ul anti-mouse ppErk1/2 antibody was added per sample in 50 ul Fc blocking solution, the cells were mixed by pipetting up and down 30x, and then incubated for 30 min at RT. The cells were washed 2x and re-suspended in 100 ul FACS buffer containing 0.1 ul Alexa Fluor Plus 555 Goat anti-mouse IgG per sample, and incubated for 20 min at RT in the dark. The cells were then washed 2x and analyzed on an Attune Nxt cytometer (Thermo Fisher Scientific).

**NFAT1 Staining**

For NFAT1 staining, cells were fixed and permeabilized *in situ* using BD Cytofix/Cytoperm kit (BD, Cat # 554714) according to the manufacturer’s protocol with some modifications. Following stimulation, half the media was carefully removed so as to not disturb the cells, and replaced with cytofix buffer and incubated for 5 min at RT. This process was repeated 2 more times with a final incubation time of 20 min at 4^o^C. Cells were gently washed 2x, the plate parfilmed, and stored at 4^o^C until all time points have been fixed. Cells were then washed 2x with 1x perm/wash buffer, and incubated for 10 min at RT after the second wash. Cells were washed 1x, stained with anti-mouse NFAT1 antibody at 1:200 in 50 ul FACS buffer and incubated for 1 hr at RT in the dark, washed 1x, stained with 0.1 ul Alexa Fluor Plus 488 Goat anti-rabbit IgG per sample in 100 ul FACS buffer and incubated for 1 hr at RT in the dark, and washed 1x. Cells were resuspended in 125 ul of a 1:1,000 dilution of DRAQ5 (Thermo Fisher Scientific, Cat # 62251) in FACS buffer and imaged using spinning disk confocal microscopy with a laser diode illuminator (AF488 - 470 Ex, 510/50 Em; DRAQ5 - 640 Ex, 700/75 Em) (89North) on the Leica DMi8 using for live imaging. Unless specified otherwise, all wash steps used FACS buffer and centrifugation at 300g for 5 min. Images were processed as described for live-imaging experiments, but with nuclear masks generated from segmented DRAQ5 images, and total cell masks generated by dilating the nuclear mask. NFAT1 fluorescence was integrated within the nuclear and total cell masks and nuclear NFAT1 was reported as the integrated nuclear signal normalized to total signal.

**Image processing and quality control**

We quantified two additional metrics for downstream quality control: (1) the fraction of H2B-iRFP signal in the nucleus, and (2) the fraction of the total mask area where the nuclear mask (M_nuc_) does not overlap the cytoplasmic mask (M_cyt_):

$$Nuclear H2B = {iRFP}_{nuc}/({iRFP}_{cyt}+{iRFP}_{nuc})$$

$$non overlapping nuclear area = Area(M_{nuc}-M_{cyt})/Area(M_{nuc}\cup M_{cyt})$$

These two metrics allow for automated exclusion of out of focus cells, dead cells, and clumped cells that would produce inaccurate pathway activity measurements (Supplementary Fig. 1B).

Following image processing, quantified cell objects were exported from MATLAB and imported into R for analysis. Individual cell objects were first filtered based on their fraction of nuclear H2B-iRFP signal (> 0.7 for inclusion) and the fraction of their nuclear mask non-overlapping with the cytoplasmic mask (< 0.1 for inclusion) (Supplementary Fig. 1B). For each of two independent live-imaging experiments, ~90k individual cell objects passed QC and proceeded to further quality control for single cell time traces. We limited our single cell time traces to 16 hrs because tracking single cells for longer than 16 hrs proved too difficult for a robust and reliable analysis. For inclusion in downstream analysis, single cell traces needed to have at least 10 measured time points. The traces that passed were then smoothed and had missing data points interpolated with local regression. Lastly, cell traces were trimmed down to 2-15 hrs due to a high prevalence of missing leading and trailing time points which cannot be interpolated, and only those cells with no missing values after this stage were included in the single cell signaling analyses. A total of 1,510 single cell traces from one experimental replicate for five pMHC conditions were used. These traces were downsampled to generate a balanced distribution for clustering analysis.

**Random forest classification models**

We first created 25 data “folds” for cross-validation using the *createMultiFolds* function from the *caret* package (v 6.0-93) in R. For each of 5, 5-fold splits, single cell traces were divided into 5 partitions while preserving the frequency of cell traces from each pMHC condition. 1 of the 5 partitions was then held out from that fold, and used for model validation. The remaining 4 partitions were used for model training. In this process, 25 independent training and validation datasets were generated such that each and every single cell trace was used to train 20 models, and held out and used for validation for the other 5 models. With these 25 data folds, we performed a nested cross-validation training and validation workflow whereby for each fold, a random forest classification model with 500 decision trees was trained and subsequently validated on the hold out partition using the *randomForest* package (v 4.7-1.1) in R. Here, each hold out partition was downsampled to create a balanced distribution of single cell traces from each pMHC condition for model validation. In this workflow, 25 independent models were trained and validated to yield 25 sets of pMHC input predictions based on various Erk and NFAT signaling predictors noted in the main text. A confusion matrix was generated from each model’s predictions, and the resulting matrices averaged to yield a final precision matrix. Overall accuracy was extracted from each model, and averaged to yield the reported model accuracies. Model accuracy p-values were generated by the *confusionMatrix* function in the *caret* package.

**Multiple linear regression models**

We used multiple linear regression modeling to estimate the effect of late signaling activity on each DEG’s expression level:

$ŷ_{gene} = \beta_{N} \cdot x_{N} + \beta_{E} \cdot x_{E} + \beta_{EN} \cdot x_{E}x_{N} + \beta_{o}$

$ŷ_{gene}$ is the log_2_ expression of a gene for a given treatment condition, measured by bulk RNA-seq, normalized by the mean expression of that gene across all treatment conditions.

$x_{N} , x_{E}$are the integrated activities of the NFAT and Erk pathways upon inhibitor treatment from 7-16 hours, measured by live imaging.

For two replicates of RNA-seq expression measurements (4 conditions x 2 replicates = 8 total data points for fitting), we fit a linear model to each of the 1,089 DEGs using the *lm* function within the *stats* package (v 4.1.2) in R with the following formula: $y \sim x_{E} * x_{N}$

**DEG tests for scRNA-seq data**

To identify pMHC affinity- and dose-dependent genes, we followed the Trapnell Lab walkthrough (<https://cole-trapnell-lab.github.io/monocle3/docs/differential/>). To identify affinity-dependent DEGs, we subsetted our cell dataset on only cells from 2 or 20 pmol M9C, or L6F stimulation, thereby comparing affinity effects at saturating doses. We used the *fit_models* function and identified DEGs using a 0.05 q-value cutoff. We did the same to identify dose-dependent DEGs, but this time subsetted our cell dataset on only cells from M9C stimulation, thereby comparing dose effects of uniform affinity pMHC. To identify stimulation-dependent, but pMHC-independent genes, we fitted models comparing all stimulation conditions to the rest condition, and identified those genes that were stimulation-dependent, but not affinity or dose dependent. We performed these DEG tests on two cell data sets: (1) the 1,089 Erk and NFAT regulated genes identified from bulk RNA-seq, and (2) all genes identified by scRNA-seq that were expressed in at least 100 cells. All of these gene lists are included in Supplementary Table 2.

**LASSO Regression**

We reasoned that genes co-expressed in single cells would be subject to the same regulatory mechanisms. To test this, we performed LASSO (least absolute shrinkage and selection operator) regression using the *glmnet* package (v 4.1-4) in R following a previously developed analysis method (63) with some modifications. First, genes expressed in fewer than 5% of the 11,412 cells in our cell dataset were excluded, leaving 810 genes. We scaled the expression of each gene as a z-score and ran the regression analysis for every pairwise set of genes. We used the pairwise regression coefficients, which estimate the correlation between two genes, to populate a gene-by-gene correlation matrix. For each pair of genes, two coefficients are generated (A ~ B and B ~ A), so we averaged the two to create a symmetrical matrix. We removed genes with no correlation to any other gene by excluding rows with a zero sum (a gene with no correlation to another gene yields a coefficient of zero), leaving 309 genes whose expression was correlated with at least one other gene. The resulting matrix was clustered using the “spearman” distance metric and the “average” hierarchical clustering method. Clusters of co-expressed genes were defined from this dendrogram by manually traversing the tree and setting cutoff heights that generated reasonable cluster boundaries based on regions of uniformly high correlation along the diagonal.

**List of Antibodies used**

PE anti-mouse CD62L (clone MEL-14; Thermo Fisher Scientific, Cat # 12-0621-82)

APC anti-mouse CD44 (clone IM7; Thermo Fisher Scientific, Cat # 17-0441-82)

BV605 anti-human CD69 (clone FN50; Biolegend, Cat # 310938)

BV605 anti-mouse CD69 (clone H1.2F3; Biolegend, Cat # 104529)

APC anti-mouse CD69 (clone H1.2F3; Biolegend, Cat # 104514)

APC anti-mouse CD25 (clone PC61.5; Thermo Fisher Scientific, Cat # 17-0251-82)

e450 anti-mouse CD279 (PD-1) (clone RMP1-30; Thermo Fisher Scientific, Cat # 48-9981-82)

Rabbit monoclonal anti-mouse NFAT1 (clone D43B1; Cell Signaling Technology, Cat # 5861)

Mouse monoclonal anti-mouse ppErk1/2 (clone 4B11B69; Biolegend, Cat # 675502)

Alexa Fluor Plus 555 Goat anti-mouse IgG (Thermo Fisher Scientific, Cat # A32727)

Alexa Fluor Plus 488 Goat anti-rabbit IgG (Thermo Fisher Scientific, Cat # A32731TR)

Ultra-LEAF Anti-mouse CD11a (Clone M17/4, Biolegend Cat # 101118)

TotalSeq^TM^-B0301 anti-mouse Hashtag 1 (clone M1/42; 30-F11; Biolegend, Cat # 155831)

TotalSeq^TM^-B0302 anti-mouse Hashtag 2 (clone M1/42; 30-F11; Biolegend, Cat # 155833)

TotalSeq^TM^-B0303 anti-mouse Hashtag 3 (clone M1/42; 30-F11; Biolegend, Cat # 155835)

TotalSeq^TM^-B0304 anti-mouse Hashtag 4 (clone M1/42; 30-F11; Biolegend, Cat # 155837)

TotalSeq^TM^-B0305 anti-mouse Hashtag 5 (clone M1/42; 30-F11; Biolegend, Cat # 155839)

TotalSeq^TM^-B0306 anti-mouse Hashtag 6 (clone M1/42; 30-F11; Biolegend, Cat # 155841)

**MATHEMATICAL APPENDIX**

**1. Modeling gene regulation to understand the relationships between Erk/NFAT signaling coefficients**

What is the basis for the strong observed negative correlation between the sensitivity coefficient for combined Erk and NFAT regulation (β_EN_) and the sensitivity coefficients for the individual pathways alone (β_E_ and β_N_)? Here, we examine the possibility that these correlations impose constraints on the *cis*-regulatory logic of target gene regulation by Erk or NFAT. To do so, we analyze a series of mathematical models of gene regulation, then derive from these models predicted relationships between these sensitivity terms, to determine whether these models can plausibly explain experimental data. First, recall that the linear model for fitting Erk and NFAT gene expression is given by:

$y(x_{N},x_{E}) = \beta_{0}+\beta_{E}{\cdot x}_{E}+\beta_{N}{\cdot x}_{N}+\beta_{EN}{\cdot x}_{E}x_{N}$	(1)

We will now construct a series of gene regulation functions for different models, incorporating different regulatory schemes for binding of NFAT or AP-1, which acts downstream of Erk activation. We will then use these gene regulation functions for different models to calculate predicted sensitivity coefficients. We will examine how sensitivity coefficients are constrained to vary with each other to see whether their modes of covariation are consistent with those observed experimentally (Fig. 4G). In these models, we make the assumption that TF binding is fast and reaches thermodynamic equilibrium (1, 2), and that each binding configuration supports a distinct transcription rate.

*1.1: Model I: Regulation by a single composite site for cooperative activation*

NFAT and AP-1 are known to bind cooperatively to composite sites on *cis*-regulatory elements, a mode of regulation known to occur for many T cell activation genes, and is therefore thought to underlie the activation of many target genes of T cell signaling. In our first model, we consider whether this mode of regulation is sufficient to account for the observed dependencies of genes on NFAT and/or Erk signaling. In this model, a single composite NFAT and AP-1 binding can be occupied jointly by both NFAT and AP-1, or singly by one factor alone. Occupancy by both factors is required for activation, though occupancy with a single factor can result in partial activation. The probabilities of finding this composite element in different transcription factor occupancy states is given by:

$\frac{dp_{N}}{dt}=\alpha_{1}n{\cdot p}_{0}-\beta_{1}{\cdot p}_{N}-\alpha_{4}a{\cdot p}_{N}+\beta_{4}p_{C}$	(2)

$\frac{dp_{A}}{dt}=\alpha_{2}a{\cdot p}_{0}-\beta_{2}{\cdot p}_{A}-\alpha_{3}n{\cdot p}_{A}+\beta_{3}p_{C}$	(3)

$\frac{dp_{C}}{dt}=\alpha_{4}a{\cdot p}_{N}-\beta_{4}{\cdot p}_{C}+\alpha_{3}n{\cdot p}_{A}-\beta_{3}p_{C}$	(4)

where $p_{N}$ , $p_{A}$ and $p_{C}$ are the probabilities of finding the element in the NFAT bound, AP-1 bound and doubly state, and $a$ and $n$ are the normalized levels of active AP-1 and NFAT transcription factors. Here, we will choose the normalization factor for these TFs such that the magnitude of these variables is unity when the corresponding factor is fully active, to enable comparison with Equation (1). Additionally, from conservation of probabilities:

$p_{0}+p_{N}+p_{A}{+p}_{C}=1.$	(5)

where $p_{0}$ is the probability of finding the element in an unbound state. Now, let the transcription rates of the gene locus in NFAT and AP-1 singly bound and doubly bound states be $v_{N}$ , $v_{A}$ and $v_{C}$. By solving equations (2)-(5), and by additionally assuming that the system is in thermodynamic equilibrium by invoking detailed balance of forward and reverse reaction rates for individual chemical binding and unbinding reactions, we get that the average rate of transcription from this gene locus as a function of NFAT and AP-1 activity levels is given by:

$v(n,a)=\frac{v_{A}a/K_{A}{+v}_{N}n/K_{N}{+v}_{C}an/K_{C}}{1+a/K_{A}+n/K_{N}+an/K_{C}}$.	(6)

where $K_{N}=\beta_{1}/\alpha_{1}$, $K_{A}=\beta_{2}/\alpha_{2}$ and $K_{C}=\beta_{2}\beta_{3}/\alpha_{2}\alpha_{3}=\beta_{1}\beta_{4}/\alpha_{1}\alpha_{4}$ are the dissociation constants for the TFs singly and in combination, in normalized units. We note that the expression for $K_{C}$ implies that a relationship between the binding and unbinding rate constants that must be satisfied for the system to be at steady state. This condition corresponds to the requirement that the system is at thermodynamic equilibrium, Now, modeling the gene expression (y) using a synthesis and first-order degradation term:

$\frac{dy}{dt}=v(n,a)-\delta y$	(7)

and hence the steady state expression level of the gene will be:

$y =v(n,a)/\delta$	(8)

Now, we consider the regime where NFAT and AP-1 bind in a highly cooperative manner such that binding of one TF enhances affinity for binding of the second TF, such that $K_{C}\ll K_{N}$, and $K_{C}\ll K_{A}$. This biochemical regime is observed with the extensive contacts between the NFAT and AP-1 upon binding to DNA (3). In this regime, by incorporating the expression for steady-state expression levels, we get:

$y_{N} = v(1,0)/\delta=\frac{v_{N}/\delta}{1+K_{N}}$	(9)

$y_{A} =$ $v(0,1)/\delta=\frac{v_{A}/\delta}{1+K_{A}}$	(10)

$y_{C}=v(1,1)/\delta=\frac{v_{C}/\delta}{1+K_{C}}$	(11)

Finally, assuming that gene is not transcribed in the absence of AP-1 or NFAT activity, we set $y_{0}=0$. Here, we note that because NFAT and AP-1 bind cooperatively ($K_{C}\ll K_{N}$, and $K_{C}\ll K_{A}$), the transcription rate when only one TF is present (Eq. 9-10) will be much lower than that when both are present (Eq. 11), and thus $y_{C}\gg y_{A}$, and $y_{C}\gg y_{N}$.

We can now solve equation (1) to get their relationship between gene expression levels and the sensitivity coefficients obtained from analysis of RNA-seq data::

$\beta_{0}{=y}_{0}$(= 0)	(12)

$\beta_{N}{=y}_{N}$	(13)

$\beta_{E}{=y}_{A}$	(14)

$\beta_{C}{=y}_{C}{-y}_{A}-y_{N}{-y}_{0}{(\sim y}_{C}\gg y_{A}, y_{N})$	(15)

From these equations, we get that these requirements translate to the condition that the sensitivity coefficients $\beta_{EN}\gg\beta_{N}$, $\beta_{EN}\gg\beta_{E}$. This is inconsistent with observations that, for most target genes, the sensitivity coefficients for single regulation ($\beta_{N}$, $\beta_{A}$) are not negligible in magnitude compared to that for joint regulation ($\beta_{C}$). A greater range of single-input sensitivities can be observed when the requirements for NFAT and AP-1 cooperativity are relaxed, such that these factors bind independently to the same composite site; however, such a binding scheme would not be consistent with the known interaction and binding between these factors observed at a biochemical and structural level.

*1.2: Model II: Regulation by multiple single and composite sites*

Our analysis of a single NFAT/AP-1 composite binding site suggest that additional elements, where NFAT or AP-1 bind either singly or combination with other factors, may be required in order to explain the observed sensitivity coefficients for target genes, where the sensitivity coefficients for single NFAT or AP-1 regulation are comparable in magnitude to that for their joint regulation, (i.e. $\beta_{N}\sim\beta_{C}$or $\beta_{A}\sim\beta_{C})$. We thus considered another *cis*-regulatory element that contains both a composite NFAT/AP-1 element, as well as two additional regulatory elements that contain binding sites for NFAT or AP-1 alone. These binding sites may either involve partnerless binding of these factors (4), or could involve binding of these factors with other partners not explicitly modeled. In any case, we can describe the binding of NFAT or AP-1 to those additional sites using additional probabilities, as follows:

$\frac{dq_{N}}{dt}=\alpha_{5}n{\cdot q}_{0}-\beta_{5}{\cdot q}_{N}$	(16)

$\frac{dr_{A}}{dt}=\alpha_{6}a{\cdot r}_{0}-\beta_{6}{\cdot r}_{A}$	(17)

where the conservation of probabilities dictates that:

$q_{0}+q_{N}=1$	(18)

$r_{0}+r_{A}=1$	(19)

Additionally, the binding of NFAT and AP-1 to the composite element is described by the same equation described above (2)-(4). Now, let us assume that each of these elements control transcription in an independent manner, such that the total average rate of transcription is given by the sum of the average transcription rates from each element (1). In such a situation, the average transcription rate from the locus is given by:

$v(n,a)=\frac{v_{C}an}{K_{C}+an}+\frac{v_{A}a}{K_{A}+a}+\frac{v_{N}n}{K_{N}+n}$	(20)

Here, we take the maximal rates of the transcription of the NFAT and AP-1 only elements to be $v_{N}$and $v_{A}$, and the dissociation constants for these TFs from their respective elements to be $K_{N}=\beta_{5}{/\alpha}_{5}$ and $K_{A}=\beta_{6}{/\alpha}_{6}$. Furthermore, we have incorporated the assumption that NFAT and AP-1 binding to the composite is highly cooperative such that there is negligible fraction of the element that is singly bound. In this regime, the expression level of the gene is given by:

$y_{N} = v(1,0)/\delta=\frac{v_{N}/\delta}{1+K_{N}}$	(21)

$y_{A} =$ $v(0,1)/\delta=\frac{v_{A}/\delta}{1+K_{A}}$	(22)

$$y_{C}=v(1,1)/\delta=\frac{v_{N}/\delta}{1+K_{N}}+\frac{v_{A}/\delta}{1+K_{A}}+\frac{v_{C}/\delta}{1+K_{C}}$$

Using equations (12)-(15), and assuming that there is no basal transcription, we then determine that the values for the sensitivity coefficients are given by:

$\beta_{0}=0$	(23)

$\beta_{N}=\frac{v_{N}/\delta}{1+K_{N}}$	(24)

$\beta_{E}=\frac{v_{A}/\delta}{1+K_{A}}$	(25)

$\beta_{C}=\frac{v_{C}/\delta}{1+K_{C}}$	(26)

With this regulatory system, we note that the maximal transcription velocities and the dissociation constants for each of the binding elements can be tuned independently, thus allowing the measured sensitivity coefficients to vary freely from each other. Such a situation can account for the variation in sensitivity coefficients between different genes, as we observed (Fig. 4); however, such a situation would be hard to reconcile with experimental observations showing the strong negative correlation between the joint sensitivity coefficient $\beta_{C}$ and the single NFAT or AP-1 sensitivity coefficients $\beta_{A}$, and $\beta_{N}$. The strength of this negative correlation and its pervasiveness across multiple gene clusters suggest the existence of additional regulatory mechanisms dictating TF occupancy and transcription rate that involve pervasive antagonism between these two transcription factors.

*1.3: Model III: Regulation by multiple sites with partner competition*

What is the basis for the observed negative correlations between the combined NFAT/AP-1 sensitivity coefficient $\beta_{C}$ and the single sensitivity coefficients $\beta_{E}$, and $\beta_{N}$? In developmental systems, it has been observed that a transcription factor can globally redirect the binding of its partner away from other sites towards the composite sites binding both TFs ^65,83^. With such a partner redirection, NFAT (or AP-1), when expressed in the cell, preferentially redirect AP-1 (or NFAT) towards composite NFAT and AP-1 elements for both TFs, at the expense of binding to elements containing only AP-1 (or NFAT) binding sites. Here, we consider a simple mathematical model for AP-1 and NFAT redirection, where these two TFs can form a heterodimer to bind the composite element:

$\frac{d[AN]}{dt}=k_{ON}[A][N]-k_{OFF}[AN]$	(27)

here, the conservation relationships for the two factors are:

$[A]+[AN] = A_{T}$	(28)

$$[N]+[AN] = N_{T}$$

where $A_{T}$ and $N_{T}$ are the total concentration of NFAT and AP-1 present when signaling is fully active. We take these quantities to be unity upon full signaling, and zero when the signaling pathway is inhibited. A full solution of (27) and (28) would involve solving a quadratic equation. Here, to gain intuition into the behavior of the system, we consider the regime where NFAT and AP-1 bind tightly to each other, and where NFAT is in abundance relative to AP-1 ($N_{T}{>A}_{T})$. In this regime, when signaling is fully active:

[A] = 0	$a=0$	(29)

[N] = $N_{T} -A_{T}$	$n=1-{(A}_{T}/N_{T})>0$	(30)

[AN] = $A_{T}$	$c=1$	(31)

where ${\rho=A}_{T}/N_{T}$ is the ratio of total AP-1 to NFAT, and where we have defined $c$ to be a variable describing the NFAT/AP-1 complex. Now, from (20), the rate of transcription is given by:

$v(n,a)=\frac{v_{C}c}{K_{C}+c}+\frac{v_{A}a}{K_{A}+a}+\frac{v_{N}n}{K_{N}+n}$	(32)

By inserting (29)-(31) into (32), we get that:

$y_{C}=v(1,1)/\delta=\frac{v_{N}(1-\rho)/\delta}{(1-\rho)+K_{N}}+\frac{v_{C}/\delta}{1+K_{C}}$	(33)

$y_{A}=v(0,1)/\delta=\frac{v_{A}/\delta}{K_{A}+1}$	(34)

$y_{N}=v(1,0)/\delta=\frac{v_{N}/\delta}{K_{N}+1}$	(35)

Using (12)-(15), and assuming negligible basal transcription, we now get that:

$\beta_{0}=0$	(36)

$\beta_{N}=\frac{v_{N}/\delta}{1+K_{N}}$	(37)

$\beta_{E}=\frac{v_{A}/\delta}{1+K_{A}}$	(38)

$\beta_{C}=\frac{v_{C}/\delta}{1+K_{C}}+\frac{v_{N}(1-\rho)/\delta}{(1-\rho)+K_{N}}-\frac{v_{N}/\delta}{K_{N}+1}-\frac{v_{A}/\delta}{K_{A}+1}$	(39)

We now note that, unlike the sensitivity coefficients derived for the multiple sites model without any partner redirection (Model II), the sensitivity coefficients for single and joint NFAT and AP-1 effects are no longer independent; specifically, when there are variations in the maximal transcription rates and dissociation constants for the NFAT and AP-1 singly bound elements ($v_{N}$, $v_{A}$, $K_{N}$, $K_{A}$), $\beta_{C}$ would now vary in the opposite direction from $\beta_{E}$ and $\beta_{N}$. For instance, in this case, amid variations in the binding parameters for the AP-1 only binding site ($v_{A}$ or $K_{A}$), we expect the following relationship:

$\beta_{C}{(v}_{A},K_{A})=C-\beta_{E}{(v}_{A},K_{A})$	(40)

where:

$C=\frac{v_{C}/\delta}{1+K_{C}}+\frac{v_{N}(1-\rho)/\delta}{(1-\rho)+K_{N}}-\frac{v_{N}/\delta}{K_{N}+1}$.	(41)

Thus, in this case, the joint sensitivity coefficient would decrease linearly with an increasing value for the AP-1 sensitivity coefficient, with an intercept given by the parameters for NFAT and composite site activity, as well as the ratios of NFAT and AP-1 levels. From these equations (36-39), we expect this inverse relationship to arise regardless of the relative strengths of the NFAT or AP-1 only site, their transcriptional rates, and we also expect this inverse relationship when the composite site is absent, such that the interaction between NFAT and AP-1 is a *trans*- effect that results in redirection of these factors towards other gene loci. These different regimes are captured by the observed regulatory modes (“A1-4” Fig. 4). By the same argument, we also expect similar inverse relationships to arise from the redirection of binding partners from the repressive binding sites. Thus, from analysis of the mathematical model, we conclude that the *trans-* redirection of NFAT and/or AP-1 binding from *cis*-regulatory elements with different strengths can plausibly explain the negative correlation between the sensitivity coefficients that is broadly observed across many different regulatory modes and clusters.

**2. Mathematical models of PD-1 and CD25 regulation**

In this section, we build a model describing the regulation of PD-1 or CD25 at the *cis*-regulatory level. We will first approximate the experimentally observed signaling dynamics using a series of simple functions. We approximate the time-dependent activity of Erk and NFAT using the following time-dependent function:

$y=b+(a-b)exp(-t/c)+d\cdot{(t/t_{end})}^{e}$	(42)

This function consists of two parts: (1) an exponential decay from an initial value *a* to a final value *b,* occurring with time constant c; and (2) a rise in activity, from an initial value of zero to a value of *d* at the end time $t_{end}$, occurring with an exponent *e*. The function captures the initial decay in signaling from a maximum value that occurs straight after signaling onset, whereas the second value captures the late rise in signaling activity that can occur with variable amplitude and sharpness. These two components of the function, together with chosen parameters, give rise to a time trace (Fig. 6E) that approximates the signaling activities observed by live cell measurements.

| Parameter | Description | Value (signaling condition) |
| --- | --- | --- |
| $a$ | initial value | 4 nM (AP-1)  9 nM (NFAT) |
| $b$ | asymptotic value of initial decay | 2 nM (AP-1)  1 nM (NFAT, L6C, 0.2 pmol M9C)  3 nM (NFAT, 2 and 20 pmol M9C) |
| c | timescale of drop off | 8 hr (AP-1)  10 hr (NFAT, L6C, 0.2 pmol M9C)  50 hr (NFAT, 2 and 20 pmol M9C) |
| $d$ | magnitude of late rise | 1 nM (AP-1, L6F)  5 nM (AP-1, M9C)  0 nM (NFAT) |
| $e$ | exponent of late rise | 3 (AP-1)  3 (NFAT) |

Table S1: Parameters used for Erk and NFAT signaling dynamics in mathematical models

The Erk and NFAT signaling levels provide a readout of the total concentration of TF in the system. Due to combinatorial action of NFAT and AP-1 transcription factors through their co-binding to common regulatory elements, there is lower free concentration of these TFs in the nucleus. To model the sequestration of free NFAT and AP-1 by binding of these factors to other elements, we utilize a simplified version of the model, where AP-1 and NFAT exist in either unbound ($A_{0},N_{0})$, singly-bound ($A_{1},N_{1})$ or doubly bound ($C$) species on DNA. Binding and unbinding reactions occur rapidly and reach equilibrium, with affinity constants that describe the initial binding of AP-1 or NFAT to DNA ($K_{N},K_{A}$), or the subsequent binding of each of these factors to form a dimeric complex on DNA ($K_{C}$):

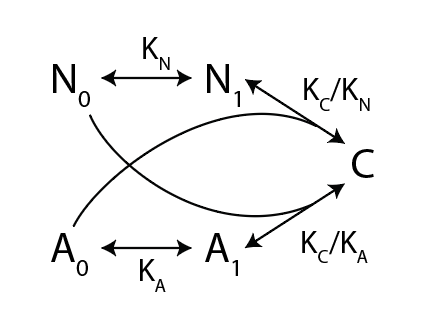

With this binding reaction scheme, it can be shown that the concentration of unbound NFAT or AP-1 that is free to act regulate the target genes is now given by to these two quadratic equations:

$K_{C}(1+K_{N})N_{0}^{2}+[(1+K_{N})(1+K_{A})+K_{C}(A_{T}-N_{T})]N_{0}-(1+K_{A})N_{T}=0$	(43)

$K_{C}(1+K_{A})A_{0}^{2}+[(1+K_{N})(1+K_{A})+K_{C}(N_{T}-A_{T})]N_{0}-(1+K_{N})N_{T}=0$	(44)

where $N_{T}$ and $A_{T}$ correspond to the total concentration of active NFAT and AP-1 factors in the nucleus as a result of signaling. For this model, we will use these equations to solve for the concentration of unbound AP-1 and NFAT given the total concentrations of these factors. These free TF concentrations along with the binding affinities of these TFs on relevant *cis*-regulatory elements at the gene locus, will then determine transcription rates for the two gene loci we consider below.

| Parameter | Description | Value (units) |
| --- | --- | --- |
| $K_{N}$ | affinity constant for NFAT to unoccupied binding site | 0.01 (1/nM) |
| $K_{A}$ | affinity constant for AP-1 to unoccupied binding site | 0.01 (1/nM) |
| $K_{C}$ | affinity constant for composite NFAT and AP-1 binding site | 0.5 (1/nM^2^) |

*2.1: Pdcd1*

From the scRNA-seq and flow cytometry data, we find that levels of *Pdcd1* transcripts, as well as PD-1 surface levels, increase in an NFAT dependent manner with both increasing affinity and dose. This suggests that a model where NFAT may bind to the *Pdcd1* locus to drive its transcription. However, because PD1 levels also increase with increasing affinity, this suggests that NFAT may function cooperatively with AP-1 factors induced by the elevated Erk signaling under stronger pMHC affinity inputs. Here, to test this hypothesis, we built a *cis*-regulatory element model of *Pdcd1* regulation, to determine whether it can plausibly explain the pMHC-dependent regulation of this gene, as mediated by NFAT and Erk/AP-1 activation. This model consists of two *cis*-regulatory elements controlling *Pdcd1* transcription: (1) a partnerless NFAT binding site; and (2) a composite NFAT/AP-1 binding site, that would be engaged in a regime with high Erk signaling levels. These two binding sites regulate *Pdcd1* transcription in an additive and independent manner, and can do so with different rates of transcription. The equation describing this mode of PD1 regulation is given by the following:

$\frac{dx}{dt}=\frac{v_{1}N_{0}}{N_{0}+K_{1}}+\frac{v_{2}(N_{0}A_{0})}{{(N}_{0}A_{0})+K_{2}}-\delta_{m}x$	(45)

$\frac{dy}{dt}=\alpha_{p}x-\delta_{p}y$	(46)

where $x$ and $y$ refer to the mRNA and protein levels for PD-1. For simulations, we modeled stochastic effects on transcription by treating the maximal transcription rates of both *cis*-regulatory elements ($v_{1},v_{2})$ as a random variable that is log-normally distributed, corresponding to the stable cell-extrinsic heterogeneity in gene expression states.

We find that the *cis*-regulatory model we describe here captures the distinct dependencies of PD-1 expression on pMHC affinity, dose and addition of signaling inhibitors. Consistent with experimental data, PD-1 expression was higher at low doses of high-affinity pMHC M9C compared to high doses of low-affinity of L6F, reflecting engagement of the NFAT:AP1 composite element due to stronger Erk signaling. PD-1 was then further up-regulated by increasing doses of M9C, reflecting increasing joint NFAT/AP-1 occupancy due to the accompanying increase in NFAT signaling. While inhibition of NFAT signaling with CsA, abolished PD-1 expression, MEKi inhibition dropped PD1 levels to those found upon stimulation with L6C, consistent with a scheme where an NFAT-only site sustains PD-1 expression in a regime where signaling activity is low. Taken together, these results are consistent with a model, where PD-1 is regulated by a bipartite non-coding element architecture, where NFAT acts singly on one site and in combination with AP-1 on the other to enable the pMHC dose and affinity specific expression of this gene.

| **Parameter** | **Description** | **Value (Units)** |
| --- | --- | --- |
| $K_{1}$ | Concentration required for half-maximal binding of NFAT to its partnerless site | 4 (nM) |
| $v_{1}$ | Maximal transcription rate induced by partnerless NFAT binding site on *Pdcd1* locus | $\mu$ = 0.2 (1/hr; mean)  ${ln(v}_{N})=N(ln(\mu),0.2)$ |
| $K_{2}$ | Concentration required for half-maximal binding of the NFAT AP-1 complex to the *cis*-regulatory site | 1900 (nM^2^) |
| $v_{2}$ | Maximal transcription rate induced by NFAT / AP-1 composite site on *Pdcd1* | 66.2 (1/hr; mean)  ${ln(v}_{N})=N(ln(\mu),0.2)$ |
| $\delta_{m}$ | mRNA degradation rate | 1 (1/hr) |
| $\alpha_{p}$ | protein synthesis rate per mRNA | $2.2\times{10}^{4}$(a.u./hr) |
| $\delta_{p}$ | protein degradation rate | 3 (1/hr) |

Table S2: Parameters for the *Pdcd1* gene regulation model.

*2.2: Il2ra*

To describe the *Il2ra* (CD25) regulation, we utilize a standard ordinary differential equation framework for gene regulation, incorporating an additional random variable to describe temporal control in the opening of this gene locus. In this model, *Il2ra* is initially in a closed state, but then transitions into an open state that can then be accessed by transcription factors for expression regulation. Based on prior studies (5, 6), we model this chromatin transition as a first-order stochastic process that occurs with a rate that is set by levels of AP-1 and NFAT transcription factors. Let $P_{1}$ be a binary random variable that is unity when the locus is open, and zero when closed. The probability:

${Pr(P}_{1}=1| t>\tau)=1-e^{-kt}$	(47)

where the first order transition rate:

$k=\frac{v_{on}(N_{0}A_{0})}{(N_{0}A_{0})+K}$	(48)

| Parameter | Description | Value (Units) |
| --- | --- | --- |
| $K$ | Concentration product required for half-maximal enhancement of *Il2ra* locus opening rate | 1.01 (nM^2^) |
| $v_{on}$ | Maximal first-order rate for *Il2ra* locus opening | 0.31 (1/hr) |
| $v_{N}$ | Maximal transcription rate induced by partnerless NFAT binding site on *Il2ra* locus | $\mu=0.7$ (1/hr; mean)  ${ln(v}_{N})=N(ln(\mu),0.2)$ |
| $K_{N}$ | Concentration for half-maximal binding / transcription | 2.7 (nM) |
| $v_{X}$ | Transcription rate due to factor X | $\mu=0.59$ (1/hr, mean, M9C)  $\mu=0.18$ (1/hr, mean, L6F)  ${ln(v}_{N})=N(ln(\mu),0.2)$ |
| $\delta_{m}$ | mRNA degradation rate | 1 (1/hr) |
| $\alpha_{p}$ | protein synthesis rate per mRNA | $2.2\times{10}^{4}$(a.u./hr) |
| $\delta_{p}$ | protein degradation rate | 3 (1/hr) |

Table S3: Parameters for the *Il2ra* gene regulation model, incorporating regulation by NFAT and AP-1 (Erk dependent).

The ordinary differential equations describing *Il2ra* regulation are given by the following variables:

$\frac{dx}{dt}=[\frac{v_{N}\cdot N_{0}}{N_{0}+K_{N}}{+v}_{X}(t)]\cdot P_{1}-\delta_{m}x$	(49)

$\frac{dy}{dt}=\alpha_{p}x-\delta_{p}y$	(50)

Here, transcription of *Il2ra* is controlled by two elements, (1) an partnerless NFAT element, which dissociation constant $K_{N}$ and maximum transcription rate $v_{N}$; and a separate element controlled by an unidentified transcription factor *X*, that drives transcription at a rate $v_{X}$. We note that this element has greater activity with high affinity pMHC (M9C) compared to with low affinity pMHC stimulation (L6F). Transcription from *Il2ra* is only possible when its locus is open, and henceforth $P_{1}=1$. One expressed, mRNA transcripts are then degraded at rate $\delta_{m}$; they are also translated at a rate of $\alpha_{p}$ per mRNA copy, and translated proteins are then degraded at a rate $\delta_{p}$. As locus opening occurs stochastically to the single-cell level, we ran multiple individual simulations of this model, each instance having a different time for locus opening dictated by the temporal probabilities, and also having varying maximal transcriptional rate by NFAT ($v_{N}$ ) and also the unidentified factor ($v_{X}$ ). These cell-to-cell variations, modeled using a log-normal distribution, provide an approximation of stable cell-to-cell heterogeneity in gene transcription due to extrinsic sources. We constrained the rate constants of the model using (1) the fractions of cells that have activated CD25 at 16 hrs (Fig. 6D), both after stimulation by the different pMHC inputs considered above and upon addition with MEKi or CsA at the onset of pMHC exposure; and (2) the mean CD25 fluorescence intensities at 30 hrs (Fig. 6D), in response to different pMHC inputs and upon inhibitor addition at 9 hrs.

This mathematical model captures both the effects of pMHC ligands, and in particular, the differential effects of signaling inhibitors on CD25 expression. In the model, early inhibition of Erk and NFAT substantially reduced the fraction of cells activating CD25, consistent with experimental observations, a consequence of disrupting NFAT or AP-1 binding at the *Il2ra* timing enhancer. In contrast, later inhibition of signaling inhibitors at 9 hrs, a time point where CD25 had already largely fully activated, resulted in much weaker effects on gene expression. NFAT inhibition moderately decreased CD25 levels, whereas MEK inhibition increased CD25 levels. These changes were explained in the model through a partnerless NFAT element whose occupancy decreased with CsA addition, but increased with MEKi addition due to redirection of NFAT away from AP-1/NFAT composite sites. We note that the reduced CD25 levels observed upon stimulation by lower-affinity L6F could not be recapitulated by the partnerless NFAT site alone, as NFAT levels are also low when the higher-affinity M9C is present at low doses. In our model, these affinity-dependent changes in expression are instead accounted for by a separate regulatory element, controlled by a separate unidentified factor X, whose activity varies with pMHC affinity.
